# A dual reporter system identifies an intermediate state and sequential regulators of 2-cell-like-to-pluripotent state transition

**DOI:** 10.1101/2021.12.17.473100

**Authors:** Chao Zhang, Jing Hao, Ming Shi, Yu-Xuan Li, Wang Yao, Yangming Wang

## Abstract

Mouse embryonic stem cells (ESCs) cycle in and out of 2-cell-like (2C-like) state in culture. The molecular mechanism governing the exit of 2C-like state remains obscure, partly due to the lack of a reporter system that can genetically mark intermediate states during exiting process. Here, we identify an intermediate state that is marked by the co-expression of <u>MER</u>V<u>L</u>::tdTomato and <u>O</u>C<u>T</u>4-GFP (MERLOT) during <u>2C</u>-like-to-<u>p</u>luripotent state transition (2CLPT). Transcriptome and epigenome analyses demonstrate that MERLOT cells cluster closely with 8-16 cell stage mouse embryos, suggesting that 2CLPT partly mimics early preimplantation development. Through a CRISPRa screen, we identify an ARRDC3-NEDD4-OCT4 regulatory axis that plays an essential role in controlling 2CLPT. Furthermore, re-evaluating previously reported 2C-like state regulators reveals dual function of Chaf1a in regulating the entry and exit of 2C-like state. Finally, ATAC-Seq footprinting analysis uncovers Klf3 as an essential transcription factor required for efficient 2CLPT. Together, our study identifies a genetically traceable intermediate state during 2CLPT and provides a valuable tool to study molecular mechanisms regulating this process.

## Introduction

Animal development is exclusively controlled by maternal factors until the activation of transcriptionally quiescent zygotic genome, a process known as zygotic genome activation (ZGA) (Lee et al., 2014). In the mouse, ZGA occurs at later 1-cell and 2-cell stages (Wu et al., 2017). During ZGA, a special group of genes (e.g. Zscan4 and Dux) (De Iaco et al., 2017; Falco et al., 2007; Hendrickson et al., 2017; Whiddon et al., 2017) and repeat sequences (e.g. murine endogenous retrovirus with leucine tRNA primer, MERVL) (Kinisu et al., 2021; Macfarlan et al., 2012; Ribet et al., 2008; Svoboda et al., 2004) are transiently expressed but quickly silenced after 4-cell stage. Coincidently, individual cells from 1-cell and 2-cell stage in the mouse have the ability to generate a full organism on their own, a capacity known as totipotency (Ishiuchi and Torres-Padilla, 2013; Lu and Zhang, 2015). The molecular mechanisms regulating the ZGA and totipotency are largely unclear due to the scarcity of cellular materials and the technical difficulty in manipulating animal embryos.

Mouse ESCs are derived from the inner cell mass of the blastocyst. These cells can self-renew indefinitely in culture and retain the ability to form three germ layers of embryo proper, a capacity known as pluripotency (Martello and Smith, 2014). In addition, mouse ESCs share some key transcriptional and functional similarities with pre-implantation epiblast, including high expression of pluripotency factors Oct4, Sox2 and Nanog and proliferation independent of ERK signaling (Boroviak et al., 2014). For these characteristics, mouse ESCs provide a unique in vitro model to understand peri-implantation development (Niwa, 2010). Interestingly, for mouse ESCs that are cultured in conventional media containing serum and leukemia inhibitory factor (LIF), a small percentage (∼0.5%) of cells are found sharing some key features with 2-cell embryos (Macfarlan et al., 2012), including the expression of 2-cell specific genes such as Zscan4 and Dux and the activation of endogenous repeat sequences such as MERVLs (De Iaco et al., 2017; Hendrickson et al., 2017; Rodriguez-Terrones et al., 2017; Whiddon et al., 2017). These cells can be genetically labeled and isolated using a MERVL long terminal repeat (LTR)-driven reporter (Macfarlan et al., 2012). In contrast to the majority of mouse ESCs, these 2C-like cells preserve higher developmental potential to differentiate into both embryonic and extraembryonic lineages (Macfarlan et al., 2012) and retain better nuclear reprogrammability (Ishiuchi et al., 2015; Iturbide and Torres-Padilla, 2020). For above reasons, studying the transition between pluripotent and 2C-like state is expected to provide insights on regulatory mechanisms underlying ZGA and totipotency.

Numerous factors, including Chaf1a/b (Ishiuchi et al., 2015), Tet2 (Guallar et al., 2018), non-canonical PRC1 complex PRC1.6 (Rodriguez-Terrones et al., 2017), Pias4 (Yan et al., 2019), miR-34 (Choi et al., 2017), Myc and Dnmt1 (Fu et al., 2019) et al. were found repressing the pluripotent to 2C-like state transition. In addition, a few other factors including Dppa2/4 (De Iaco et al., 2019; Eckersley-Maslin et al., 2019; Yan et al., 2019), Dux (De Iaco et al., 2017; Fu et al., 2019; Hendrickson et al., 2017; Whiddon et al., 2017) and Nelfa (Hu et al., 2020) were found promoting the pluripotent to 2C-like state transition. In contrast to pluripotent to 2C-like state transition, much less is known about factors controlling 2CLPT. In a recent report, Fu et al. identify an intermediate state during 2CLPT through transcriptomic analysis (Fu et al., 2020). They further discover Smg7 and nonsense mediated decay (NMD) promoting 2CLPT, possibly through destabilizing Dux mRNA. The discovery of the intermediate state in this study offers a first glance on the regulation of 2CLPT. However, the lack of appropriate genetic marker prevents the isolation and further characterization of the intermediate state. Therefore, identification of a genetically marked intermediate state will likely provide more insights on the regulation of 2CLPT.

The transcription factor *Oct4* (also known as *Pou5f1*) plays important roles in the maintenance of pluripotent state and reprogramming (Jaenisch and Young, 2008; Nichols et al., 1998; Radzisheuskaya and Silva, 2014; Shi and Jin, 2010; Takahashi and Yamanaka, 2006). Interestingly but perhaps not surprisingly, the OCT4 protein is reported absent in 2C-like cells which are considered in a totipotent-like instead of pluripotent state (Ishiuchi et al., 2015; Macfarlan et al., 2012). In this study, we identify a rare population of cells marked by concurrent expression of OCT4 protein and MERVL repeats. These cells can be genetically labeled using OCT4-GFP fusion protein and MERVL-LTR driven tdTomato reporters. We dub <u>MER</u>V<u>L</u> and <u>O</u>C<u>T</u>4 double positive cells as MERLOT cells. We demonstrate that MERLOT cells represent the intermediate state during 2CLPT by cell fate tracing through real-time microscopic imaging and flow cytometry analyses. Transcriptome and epigenome analyses further support that MERLOT cells are intermediates between 2C-like and pluripotent cells. Interestingly, MERLOT cells cluster closely with 8-16 cell stage embryos at transcriptome and epigenome levels, indicating that 2CLPT may partly mimic early preimplantation development. Through a CRISPR screen for regulators in protein degradation pathway, we find that OCT4 protein is degraded by ARRDC3-NEDD4-proteasome pathway during the entry of 2C-like state and its re-stabilization in MERLOT cells promotes the exit of 2C-like state. In addition, we uncover Chaf1a, Klf3, Smg7 and Upf1 as important regulators promoting 2CLPT at different steps. Together, our study identifies a genetically labeled and isolatable intermediate state during 2CLPT and provides a valuable tool to study molecular mechanisms regulating 2CLPT.

## Results

### OCT4 protein is expressed in a subset of 2C::tdTomato^+^ ESCs

We first checked OCT4 protein level in 2C::tdTomato reporter ESCs (Macfarlan et al., 2012; Yan et al., 2019) by immunofluorescence staining (IF). Consistent with previous studies (Ishiuchi et al., 2015; Macfarlan et al., 2012), we found that OCT4 protein is absent in most 2C::tdTomato^+^ ESCs (**Figure S1A**). Interestingly, we also noticed that some 2C::tdTomato^+^ ESCs express a relatively high level of OCT4 protein (**Figure S1A**). Flow cytometry analysis showed that 0.27% and 0.49% 2C::tdTomato^+^ cells are OCT4 positive and negative (**Figure S1B**), respectively. In addition, 3.83% 2C::tdTomato^-^ cells were OCT4^-^ (**Figure S1B**). Dux is known to promote 2C-like transition (De Iaco et al., 2017; Fu et al., 2020; Fu et al., 2019; Hendrickson et al., 2017; Whiddon et al., 2017). We constructed a doxycycline-inducible Dux transgene in 2C::tdTomato reporter ESCs (**Figure 1A**). Consistent with previous studies (Fu et al., 2020; Fu et al., 2019), the percentage of 2C::tdTomato^+^ cells is evidently increased upon transient Dux induction (**Figure S1B**). In this case, we observed around one fourth of 2C::tdTomato^+^ cells expressing OCT4 protein (**Figure S1B**). Interestingly, we also noticed that the fraction of 2C::tdTomato^-^OCT4^-^ cells is markedly increased upon Dux induction (**Figure S1B**). Together, these data demonstrate that OCT4 protein is expressed in a subset of 2C::tdTomato^+^ cells.

**Figure 1:**
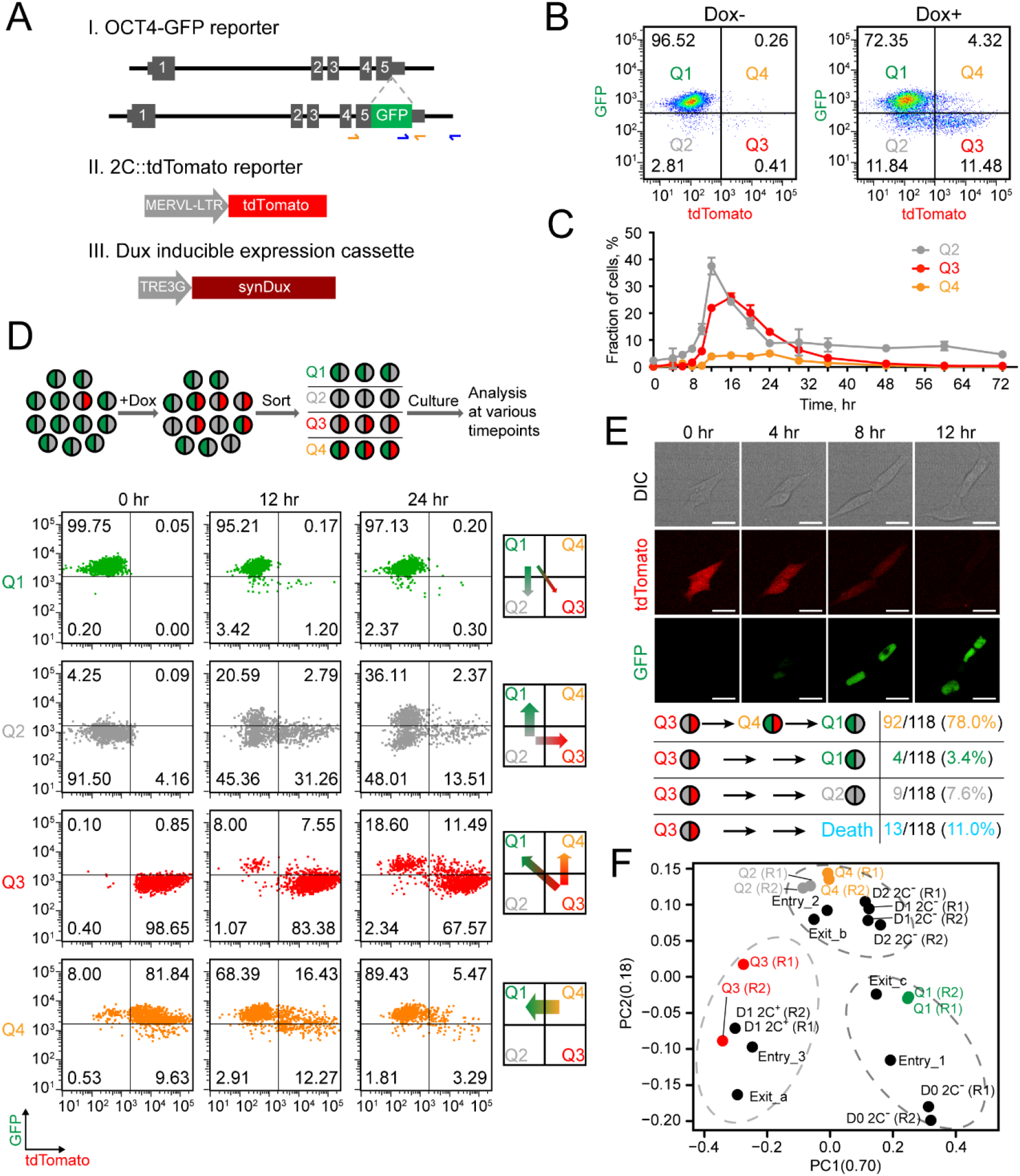
MERLOT cells represent an intermediate state during 2CLPT. (A) Design of 2C::tdTomato and OCT4-GFP reporters and inducible expression cassette of Dux. (B) Flow cytometry analysis of 2C::tdTomato/OCT4-GFP double reporter ESCs with or without Dux induction by 40 nM doxycycline. The fractions of Q1/2/3/4 are indicated. (C) Fraction of Q2/3/4 upon Dux induction by 40 nM doxycycline over time. Shown are mean ± SD, n = 3 independent experiments. (D) Dynamic transition of Q1/2/3/4 over time. Upper, experimental design; Lower and left, flow cytometry analysis of sorted Q1/2/3/4 at 0, 12 and 24 h; Lower and right, cartoon model of dynamic transition of Q1/2/3/4. This experiment was performed independently four times with similar results. (E) Live imaging of dynamic transition of Q3. Upper, representative images of dynamic transition of Q3, scale bar, 20 µm; Lower, the fraction of Q3 in different transition path. Results shown are pooled from 3 independent cultures. 118 cells with tdTomato signals diminished were captured and traced by time-lapse confocal microscope. (F) Transcriptome-based PCA of Q1/2/3/4 and intermediate states defined by Fu and coauthors (Fu et al., 2019; Fu et al., 2020).

### OCT4-GFP reporter faithfully reflects the level of OCT4 protein in ESCs

To genetically label 2C::tdTomato^+^OCT4^+^ cells, we knocked in a green fluorescent protein (GFP) to the carboxyl terminus of OCT4 protein at one allele so that this locus will produce OCT4-GFP fusion protein while the other locus produces wild type OCT4 protein (**Figures 1A** **and S1C**). OCT4 antibody staining followed by flow cytometry analysis showed that OCT4-GFP reporter faithfully reflected the level of OCT4 protein in individual ESCs (**Figure S1D**). Consistent with results from OCT4 antibody staining (**Figure S1B**), we observed 96.52% 2C::tdTomato^-^OCT4-GFP^+^ (Q1), 2.81% 2C::tdTomato^-^OCT4-GFP^-^ (Q2), 0.41% 2C::tdTomato^+^OCT4-GFP^-^, 0.26% 2C::tdTomato^+^OCT4-GFP^+^ cells (**Figure 1B**). Upon Dux induction, we observed around one fourth 2C::tdTomato^+^ cells expressing OCT4-GFP (**Figure 1B**). To figure out how OCT4 protein is repressed, we constructed OCT4-T2A-GFP reporter ESCs in which a T2A peptide (Doronina et al., 2008) that induces ribosome skipping is inserted between OCT4 and GFP (**Figure S1E**), therefore decoupling the stability control of OCT4 and GFP. Compared to OCT4-GFP reporter ESCs, GFP^-^ cells were mostly diminished in OCT4-T2A-GFP reporter ESCs. These data suggest that OCT4 protein is repressed majorly through the protein degradation pathway. However, some GFP^-^ cells were still present in OCT4-T2A-GFP reporter ESC culture (**Figure S1E**), suggesting the existence of translational and/or transcriptional repression of OCT4 protein.

### 2C-like cells become MERLOT cells during 2CLPT

To figure out the sequential order of the generation of Q2, Q3 and Q4 during Dux induction, we performed flow cytometry analysis of 2C::tdTomato/OCT4-GFP reporter ESCs at various timepoints after transient induction of Dux. We found that the fraction of Q2 cells reach the peak first, followed by Q3 and Q4 (**Figure 1C**). These results suggest a sequential transition order of Q1 to Q2 to Q3 to Q4. To further clarify this sequential order, we sorted out Q1/2/3/4 cells, re-plated them in serum/LIF media and then analyzed them by flow cytometry after culturing for 12 and 24 hours (**Figure 1D**). We found that Q2 produces Q3, Q3 produces both Q4 and Q1, while Q4 majorly produces Q1. We suspect that a large fraction of Q1 generated from Q3 may go through Q4 stage first since the rate of Q4 to Q1 conversion is much faster than Q3 to Q4 (**Figures S2A and S2B**). To clarify this, we sorted out Q3 and tracked the change of GFP and tdTomato by real-time microscopy imaging (**Figure 1E**). We found that direct conversion of Q3 to Q1 is minimum and the majority of Q3 follow a path of Q3 to Q4 to Q1. These data strongly suggest that Q4 (MERLOT cells) represent an intermediate state of 2CLPT. Moreover, kinetics analysis suggest that 2C-like (Q3) to MERLOT (Q4) state transition is the rate limiting step during the exit of 2C-like state (**Figure S2B**).

### MERLOT cells show an intermediate profile between 2C-like and pluripotent ESCs at transcriptome and epigenome levels

Next we analyzed the transcriptome of Q1/2/3/4 cells by RNA-Seq. We performed principal component analysis (PCA) on these cells along with RNA-Seq data from Yi Zhang’s group (Fu et al., 2020; Fu et al., 2019). As expected, Q1 and Q3 clustered closely with pluripotent cells and 2C-like cells (**Figure 1F**), respectively. Consistent with flow cytometry and microscopy imaging analysis, Q4 (MERLOT) clustered closely with cells designated as intermediates (Fu et al., 2020; Fu et al., 2019) between 2C-like and pluripotent ESCs (**Figure 1F**). D2 2C^-^ cells (Fu et al., 2020) represent an intermediate state identified by Fu et al. during the exit of 2C-like state. Because 2C::tdTomato is silenced in these cells, we conjecture that MERLOT represents a different intermediate state earlier than D2 2C^-^ cells. Consistent with this hypothesis, pluripotency genes were lower and 2C genes were higher in MERLOT than D2 2C^-^ cells (**Figures S2C and S2D**). In addition, we found that Q2 also clusters closely with other intermediates (**Figure 1F**). Along with flow cytometry analysis showing that Q2 cells give rise to Q3 cells (**Figure 1D**), these data suggest that Q2 may represent an intermediate state during the entry of 2C-like state. Together, transcriptome analysis supported that MERLOT cells represent an intermediate state during the exit of 2C-like state.

We then characterized expression change of protein coding genes in Q1/2/3/4 and identified four major class of genes which exhibit different dynamics (**Figures 2A and 2B**). Among them, group 1 genes (n=5368) decreased during pluripotent to 2C-like state transition while increased during the exit of 2C-like state (**Figures 2A and 2B**); many pluripotency genes and genes in stem cell maintenance, purine and glutathione metabolism pathways are enriched in group 1 (**Figures 2C and 2D**). Group 2 genes are relatively enriched in Q2 which included many 4-cell embryo activated (4C) genes and are enriched in many signaling pathways including AMPK and MAPK (**Figures 2A-2D**). Contrary to group 1 genes, group 3 genes (n=1253) increased during pluripotent to 2C-like state transition while decreased during the exit of 2C-like state. As expected, 2-cell embryo specific (2C) genes are enriched in group 3 (**Figure 2C**). In addition, genes in p53, MAPK and FOXO signaling pathways were also found enriched in group 3 (**Figure 2D**). Finally, group 4 genes (n=107) show continuous upregulation pattern during Q2 to Q3 and Q3 to Q4 transition (**Figures 2A and 2B**). Notably, group 4 included some 8-cell embryo activated (8C) genes and enriched genes in Wnt signaling and ribosome pathways (**Figures 2C and 2D**).

**Figure 2:**
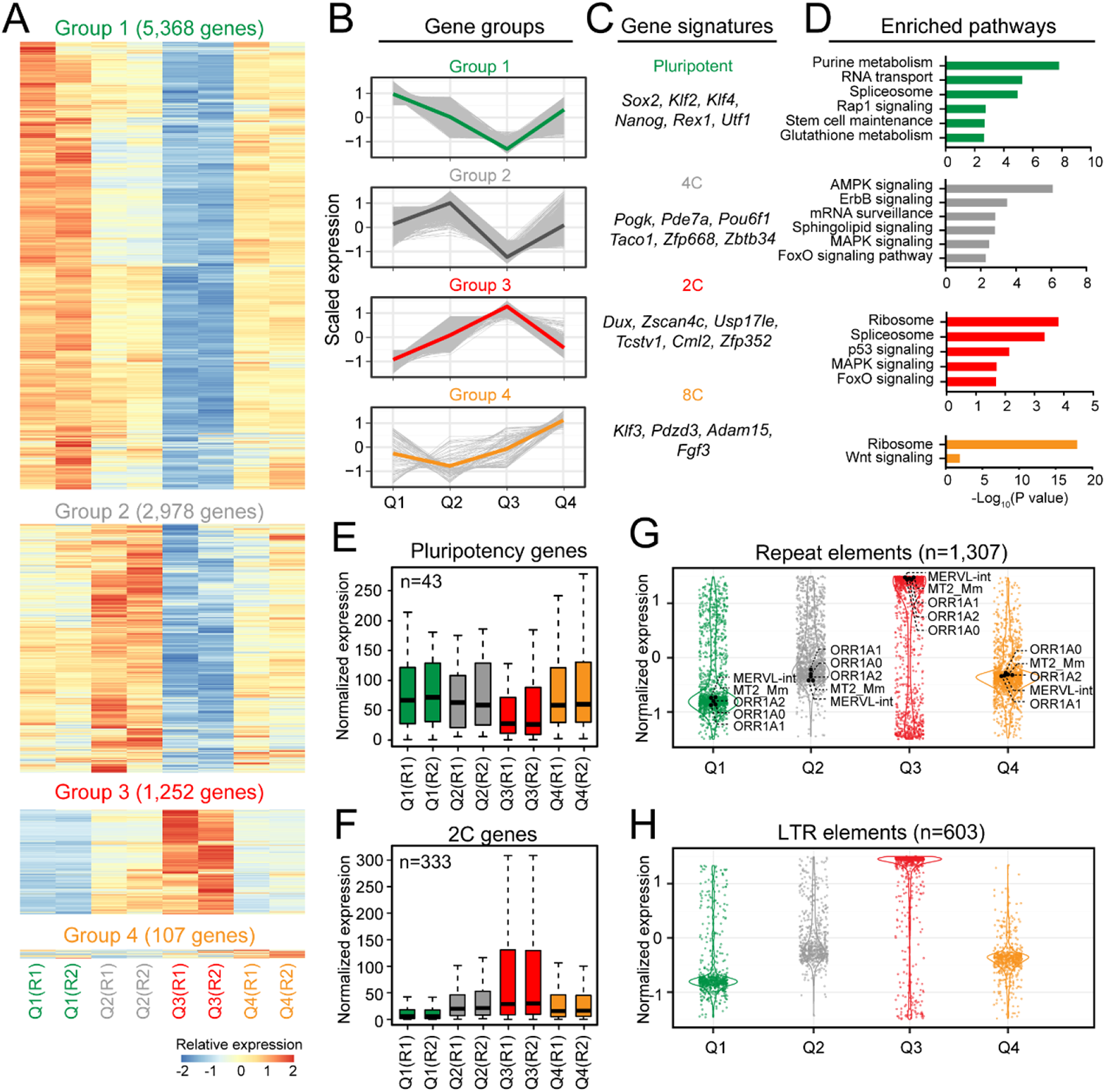
MERLOT cells display intermediate expression of pluripotency and 2C-specific genes. (A) Heatmap showing the relative expression level of 4 groups of genes defined by expression dynamics in Q1/2/3/4. Data were from two biologically independent samples of Q1/2/3/4. (B) Line graph showing the expression dynamics of group 1 to 4 genes in Q1/2/3/4. Each gene is shown as a gray line; the colored line represents the mean expression. (C) Gene signatures in each gene group. (D) The enrichment of KEGG pathway in each gene group. Selected pathways are shown and ranked according to -log_10_ (*p*-value) value. (E and F) Box and whisker plot showing the expression level of (E) pluripotency genes (n=43) and (F) 2C specific genes (n=333) in Q1/2/3/4 from two biologically independent samples. Center line, median; Box limits, upper and lower quartiles; Whiskers, 1.5× interquartile range. (G and H) Violin plot showing the relative expression of (G) all repeat elements (n=1,307) and (H) LTR elements (n=603) in Q1/2/3/4. MERVL-int, MT2_Mm, ORR1A0/1/2 are indicated. Each dot represents one repeat element. Shown are average expression of two biologically independent samples.

We then focused our analysis on pluripotency genes and 2C specific genes, signatures of pluripotent and 2C-like state. Overall, the expression of these signature genes in Q4 and Q2 displayed an intermediate level between 2C-like and pluripotent state (**Figures 2E and 2F**), again supporting that Q2 and Q4 are the intermediate state during the entry and exit of 2C-like state. Consistent with these results, the expression of repeat elements, in particular LTR elements, are generally upregulated in 2C-like state compared to pluripotent state, and are downregulated in Q4 during the exit of 2C-like state (**Figures 2G and 2H**). Together, these results show that MERLOT cells display an intermediate profile between 2C-like and pluripotent ESCs at transcriptome level.

Next we analyzed chromatin accessibility by assay for transposase-accessible chromatin (ATAC) sequencing (ATAC-seq) (Buenrostro et al., 2013), histone 3 lysine 9 trimethylation (H3K9me3) and histone 3 lysine 27 acetylation (H3K27ac) by CUT&Tag (Kaya-Okur et al., 2019) in Q1/2/3/4. ATAC and H3K27ac signals correlate with gene activation, while H3K9me3 signals correlate with gene repression. Consistent with expression changes, ATAC and H3K27ac signals of pluripotency genes were downregulated (**Figures 3A and 3B**), while ATAC and H3K27ac signals of 2C specific genes were upregulated in Q3 versus Q1 (**Figures 3C and 3D**). In contrast, H3K9me3 signals of pluripotency genes were generally very low and largely unchanged while those of 2C specific genes were downregulated in Q3 versus Q1 (**Figures 3E and 3F**). Furthermore, similar to those of 2C specific genes, ATAC and H3K27ac signals of MERVL were upregulated (**Figures S2E and S2F**), and H3K9me3 signals of MERVL were downregulated in Q3 versus Q1 (**Figure S2G**). In most cases, we observed an intermediate profile of Q4 and Q2 between Q3 and Q1 (**Figures 3A-3G and S2E-S2G**). Altogether, these data show that MERLOT cells have an intermediate epigenome profile between 2C-like and pluripotent ESCs.

**Figure 3:**
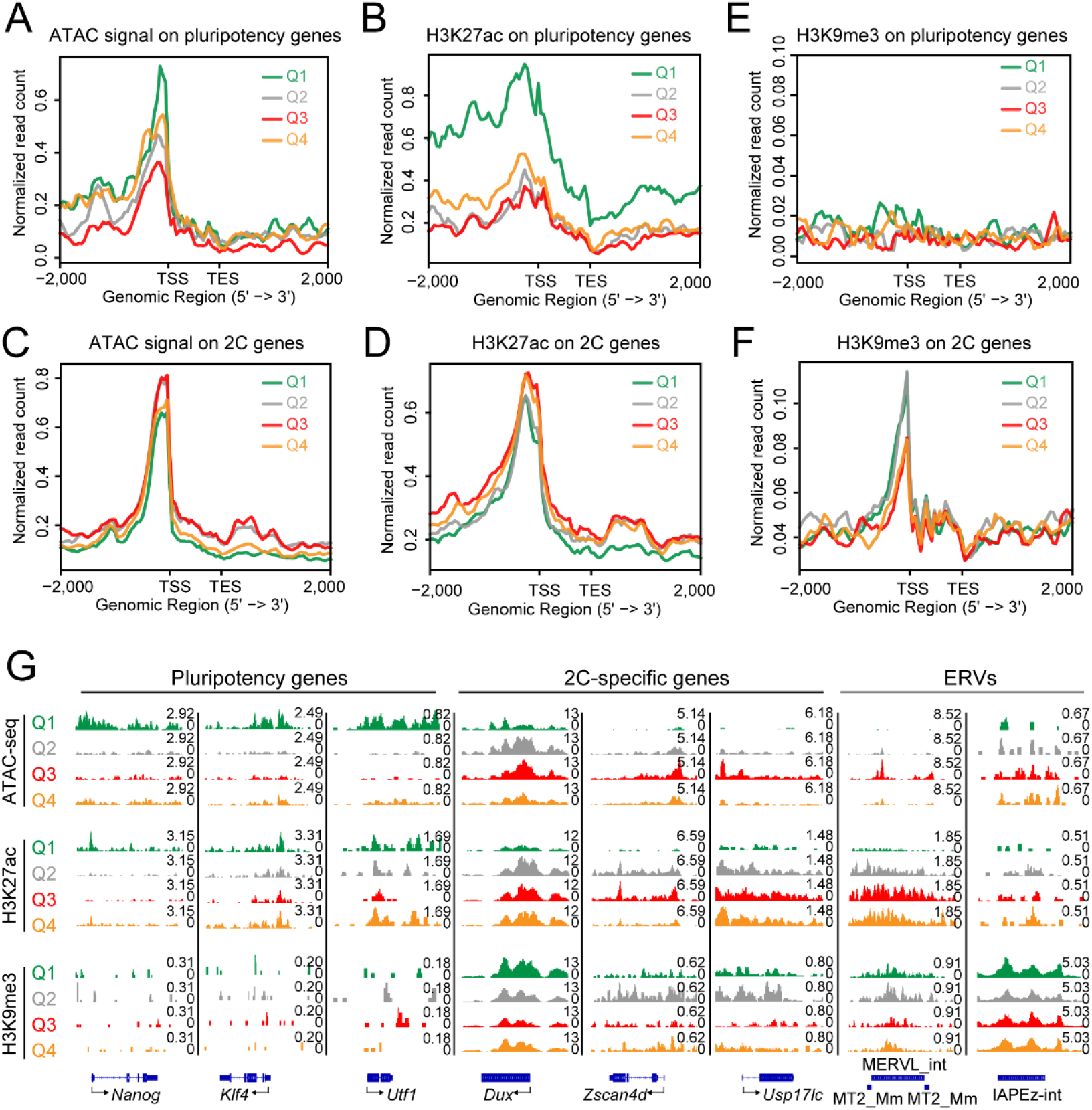
MERLOT cells display intermediate epigenome profiles on pluripotency and 2C-specific genes. (A and B) Read-count tag density pileups of (A) ATAC signal and (B) H3K27ac profile on pluripotency genes (n=43). TSS, transcription start site; TES, transcription end site. (C and D) Read-count tag density pileups of (C) ATAC signal and (D) H3K27ac profile on 2C specific genes (n=333). (E and F) Read-count tag density pileups of H3K9me3 profile on (E) pluripotency genes (n=43) and (F) 2C specific genes (n=333). (G) Genomic views of ATAC, H3K27ac and H3K9me3 CUT&Tag signals on representative pluripotency genes (*Nanog*, *Klf4*, *Utf1*), 2C specific genes (*Dux*, *Zscan4d*, *Usp17lc*) and ERVs (MERVL, IAP). For all panels, shown are the average of two biologically independent samples.

### MERLOT cells share similarities with 8-16 cell stage embryos transcriptionally and epigenetically

As 2C-like cells and ESCs exhibit similar features with 2-cell stage and blastocyst embryo, respectively, we wondered whether 2CLPT could recapitulate some of features during early embryonic development. Interestingly, genes enriched in Q4 were found activated at 8-cell stage embryos (Deng et al., 2014) (**Figure 4A**). Furthermore, PCA analysis of RNA-Seq data showed that Q4 cells cluster closely with 8-cell and 16-cell embryos (**Figure 4B**). Consistently, 8-cell and 16-cell embryo specific genes were highly enriched in Q4 versus Q3 (**Figures 4C and 4D**). Furthermore, ATAC-Seq showed that gained and lost signals in 8-cell embryos versus 4-cell embryos (Wu et al., 2016) are also gained and lost in Q4 versus Q3 (**Figure 4E**), respectively. Together, these data demonstrate that the transcriptional and epigenetic profiles of MERLOT cells share some similarities with those of 8 and 16-cell stage embryos.

**Figure 4:**
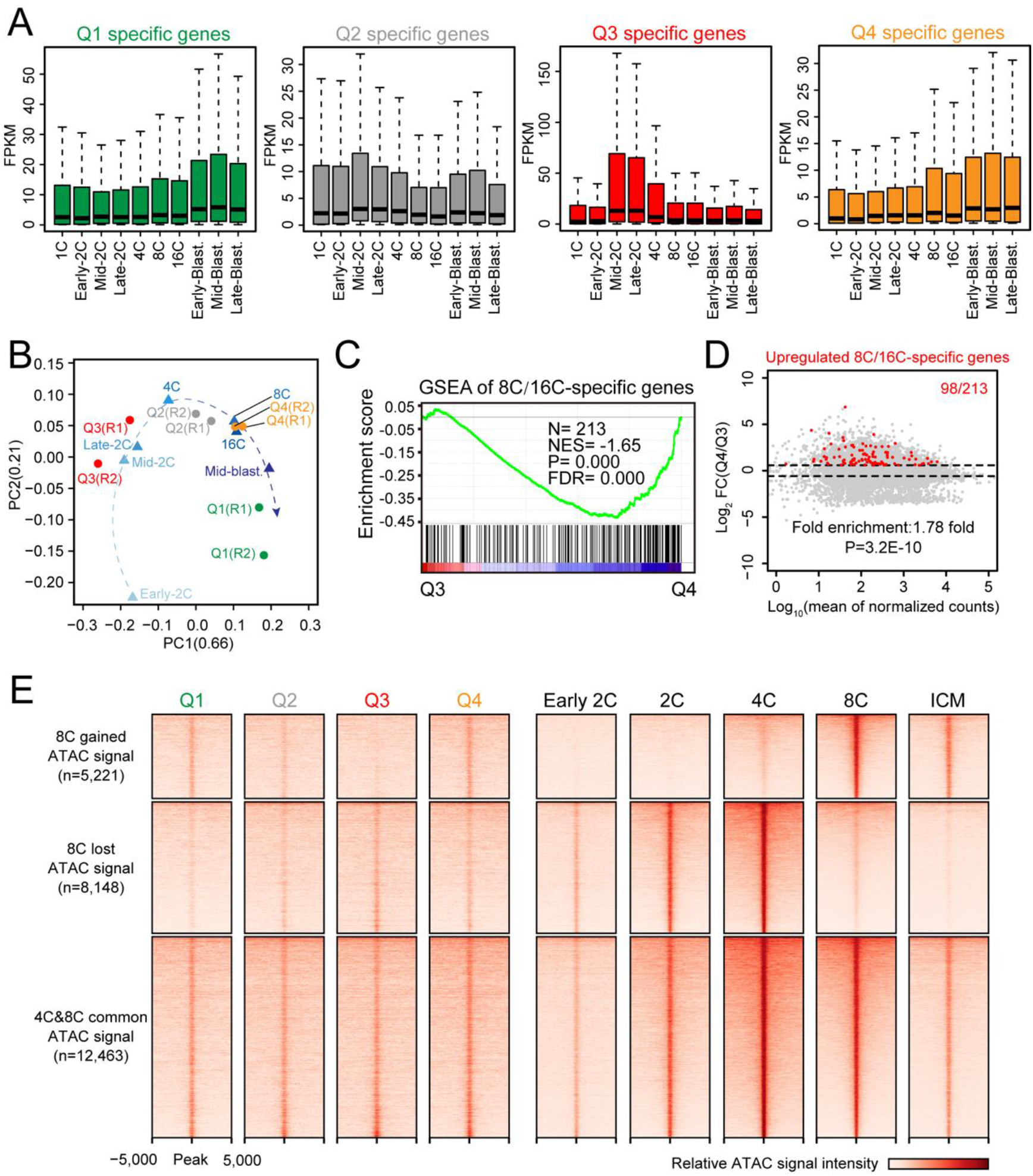
MERLOT cells share similar transcriptome and chromatin accessibility profiles with 8 and 16 cell stage embryos. (A) Box and whisker plots showing the expression of Q1/2/3/4 specific genes in mouse preimplantation embryos. Center line, median; Box limits, upper and lower quartiles; Whiskers, 1.5× interquartile range. (B) Transcriptome-based PCA of Q1/2/3/4 and mouse preimplantation embryos. (C) GSEA for 8C/16C embryo-specific genes in Q3 and Q4. (D) MA plot showing gene expression changes in Q4 versus Q3. 8C/16C embryo-specific genes are indicated by red dots. The *p*-value was calculated by hypergeometric test. (E) Heat map showing regions of ATAC signal gained, lost and unchanged in 8C versus 4C embryos in Q1/2/3/4 and early embryos. For (A) and (B), RNA-Seq data of pre-implantation embryos are from Deng et al.’s study (Deng et al., 2014). For (E), ATAC-Seq data of preimplantation embryos are from Wu et al.’s study (Wu et al., 2016).

### The re-stabilization of OCT4 promotes the exit of 2C-like state

To gain insights on the regulatory mechanism behind the exit of 2C-like state, we first checked the functional role of OCT4 in 2CLPT. Consistent with protein expression of OCT4, footprinting analysis of ATAC-Seq data using TOBIAS (Bentsen et al., 2020) revealed that OCT4 signals were downregulated during pluripotent to 2C-like state transition and were reverted in MERLOT cells during the exit of 2C-like state (**Figure 5A**). In addition, SOX2 showed a similar pattern of footprinting signal changes as OCT4 (**Figure 5A**). We then sorted out Q2/3/4 cells and transfected them with synthetic Oct4 and Sox2 mRNA. Oct4 but not Sox2 mRNA significantly promoted the exit of 2C-like state (Q3) (**Figures 5B and 5C**). Oct4^5R^ carrying 5 lysine residues (K) substituted with arginine residues (R) is resistant to proteasome degradation (Li et al., 2018). As expected, Oct4^5R^ produced higher effect than Oct4 in promoting the exit of 2C-like state (**Figures 5B and 5C**). We noticed that Oct4 and Oct4^5R^ also promote the reversion of Q2 back to Q1 and slightly block the entry of Q2 into 2C-like state (**Figures 5B and 5C**). Furthermore, Oct4 and Oct4^5R^ slightly promoted Q4 to Q1 transition (**Figures 5B and 5C**). These data suggest that OCT4 mainly promotes 2C-like to MERLOT state transition, the rate limiting step during the exit of 2C-like state.

**Figure 5:**
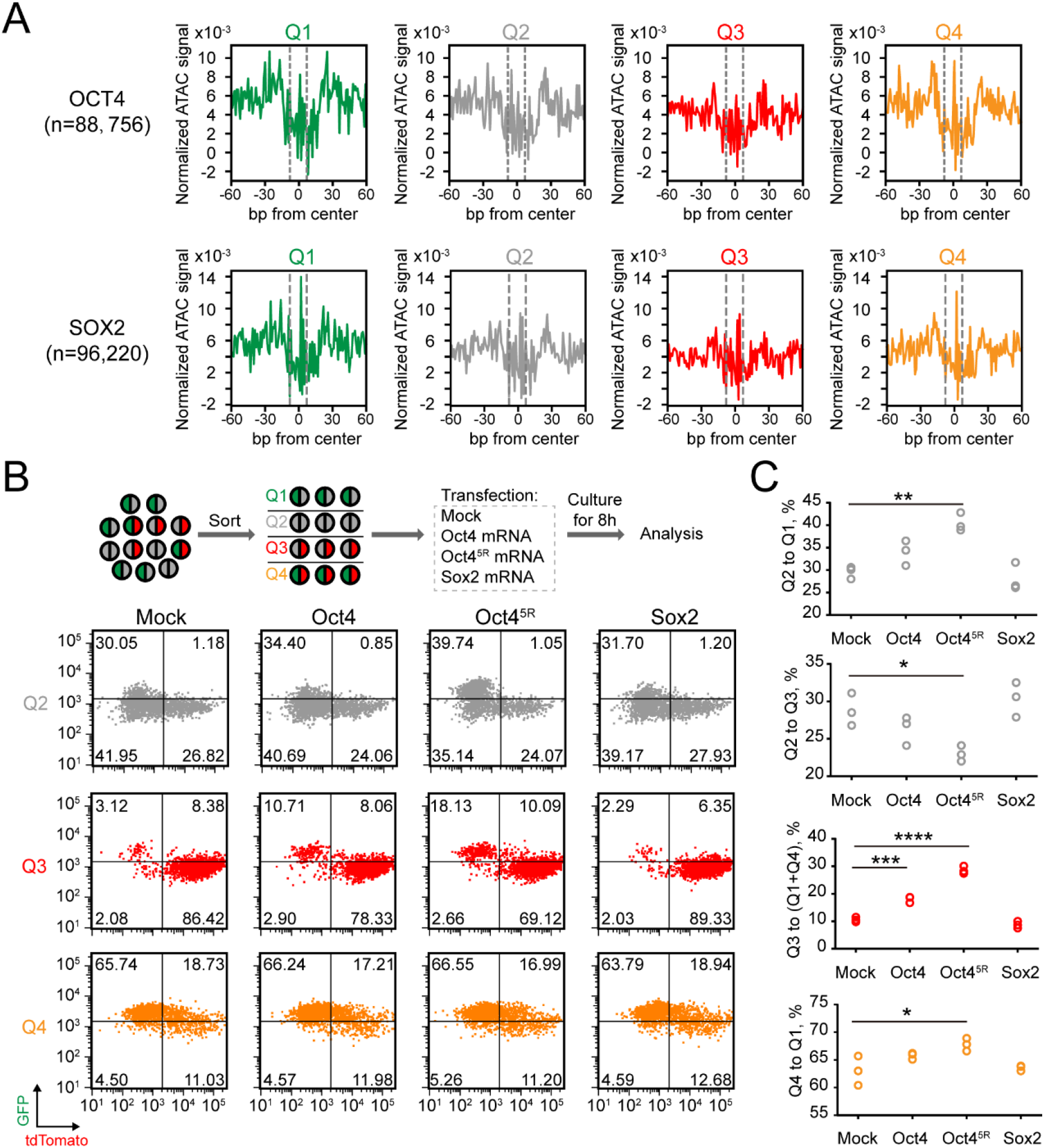
The re-stabilization of OCT4 promotes 2CLPT. (A) Aggregated footprint plots for OCT4, SOX2 in Q1/2/3/4. The dashed lines represent the edges of TF motif. (B) Flow cytometry analysis of dynamic transition of sorted Q2/3/4 transfected with Oct4, Oct4^5R^ and Sox2 mRNA. Upper, experimental design; Lower, representative dot plot. (C) Quantification of the percentage of indicated populations from (B); each circle represents one independent experiment, n = 3. The *p*-value was calculated by one-way ANOVA followed by two-tailed Dunnett’s test.

### OCT4 protein is degraded through ARRDC3-NEDD4-proteasome pathway

Next we investigated how OCT4 protein is repressed during pluripotent to 2C-like state transition. Upon Dux induction, OCT4 protein was significantly decreased as the fraction of 2C-like cells increased (**Figures S3A-S3E**). In contrast, Oct4 mRNA level was largely unchanged (**Figure S3F**). Addition of proteasome inhibitor MG132 significantly rescued OCT4 protein level (**Figures S3D-S3F**). Furthermore, Dux induction evidently increased the ubiquitination level of OCT4 which was further increased upon the addition of MG132 (**Figure S3G**). These data demonstrate that OCT4 protein is degraded through ubiquitin-proteasome pathway during the entry of 2C-like state.

To identify the pathway responsible for OCT4 degradation, we focused on ubiquitin-proteasome pathway related genes that are upregulated in both Q2 and Q3 (Q2_TPM_/Q1_TPM_>1.3, Q3_TPM_/Q1_TPM_>2, Q2_TPM_>10, Q3_TPM_>10), as well as three E3 ligases (WWP2, ITCH, TRIM32) (Bahnassawy et al., 2015; Liao et al., 2013; Xu et al., 2004) that have been reported to regulate OCT4 degradation **(Figure S4A)**. We then performed CRISPRa screening for these genes in 2C::tdTomato/OCT4-GFP reporter ESCs (**Figure S4B**). The results showed that activating Arrdc3, Arrdc4, Fbxo15 and Lonrf3 significantly decreased the OCT4-GFP signal (**Figure 6A**). The function of Arrdc3 and Arrdc4 in decreasing OCT4-GFP signal was further verified by the induction of a transgene (**Figures S4C and S4D**). Furthermore, deletion of *Arrdc3* but not *Arrdc4* rescued OCT4 protein level and decreased the expression of MERVL upon Dux induction (**Figures S5A-S5D**), suggesting that Arrdc3 is majorly responsible for OCT4 protein degradation during the induction of 2C-like state. Arrdc3 is an ubiquitin E3 ligase adaptor of α-arrestin family that typically recruits NEDD4 family of E3 ligases (Nabhan et al., 2010). Immunoprecipitation with OCT4 antibody supported the direct interaction between OCT4 and ARRDC3 (**Figure S5E**). Western blotting analysis confirmed that ARRDC3 promotes the degradation of OCT4 protein through the ubiquitin-proteasome pathway (**Figures 6B and 6C**). Furthermore, knocking down Nedd4 but not other NEDD4 family E3 ligases suppressed Arrdc3-mediated downregulation of OCT4-GFP (**Figures S5F-S5H**). Finally, mutation of PY domain which is essential for the recruitment of ubiquitin E3 ligases (Nabhan et al., 2010) in ARRDC3 diminished its ability to promote the degradation and ubiquitination of OCT4 (**Figures 6B and 6C**). Together, these data suggest that OCT4 is repressed by ARRDC3-NEDD4-proteasome pathway during pluripotent to 2C-like state transition.

**Figure 6:**
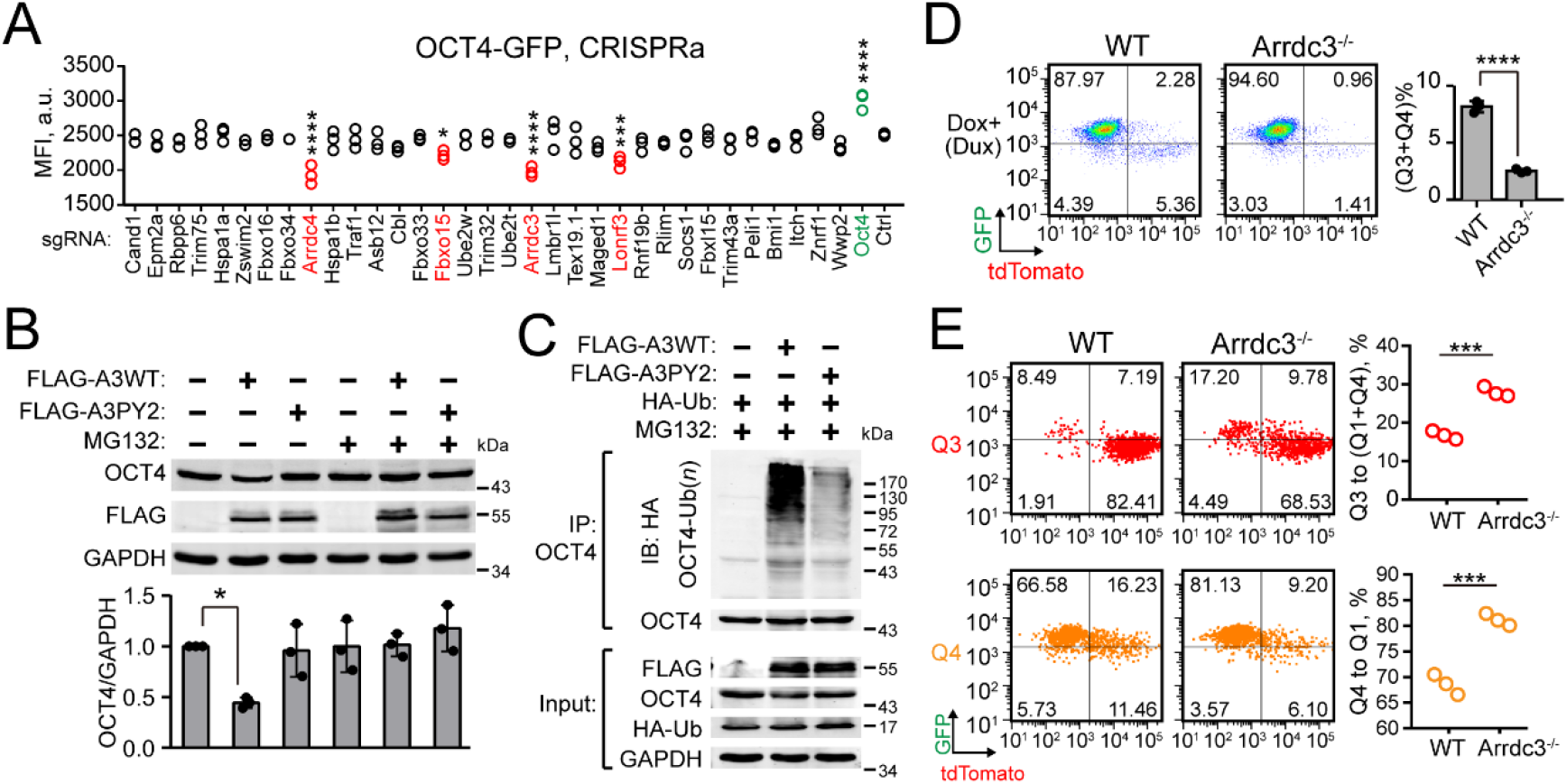
ARRDC3 promotes the degradation of OCT4 through proteasome pathway. (A) Quantification of fluorescence intensity of OCT4-GFP in CRISPRa screening. Positive candidates are labeled in red; Oct4 sgRNA as positive control is labeled in green. Each circle represents one independent experiment, n = 3. The *p*-value was calculated by one-way ANOVA followed by two-tailed Dunnett’s test. (B) Western blot analysis of OCT4 protein upon Arrdc3 or Arrdc3 PY2 mutant overexpression with or without MG132 treatment. Upper, representative images of western blot; Lower, quantification of OCT4 protein. Shown are mean ± SD, n = 3 independent experiments. The *p*-value was calculated by one-way ANOVA followed by two-tailed Dunnett’s test. (C) Western blot analysis of OCT4 ubiquitination in OCT4 immunoprecipitated samples upon Arrdc3 or Arrdc3 PY2 mutant overexpression. (D) Flow cytometry analysis of wild-type or Arrdc3 knockout ESCs with or without Dux induction by 20 nM doxycycline. Left, representative dot plots; Right, quantification of the percentage of Q3+Q4. Shown are mean ± SD, n = 3 independent experiments. The *p*-value was calculated by unpaired two-tailed Student’s *t* test. (E) Flow cytometry analysis of dynamic transition of sorted Q3 and Q4 from wild-type or Arrdc3 knockout ESCs. Left, representative dot plots; Right, quantification of the percentage of indicated populations; each circle represents one independent experiment, n = 3. The *p*-value was calculated by unpaired two-tailed Student’s *t* test.

We then checked the fraction of 2C-like cells in *Arrdc3*^-/-^ ESCs upon the induction of Dux. Consistently, we observed significant reduction of 2C-like cells in *Arrdc3*^-/-^ ESCs (**Figure 6D**). In addition, the rate of exiting 2C-like state is significantly increased in *Arrdc3*^-/-^ ESCs (**Figure 6E**). Together, these data demonstrate that ARRDC3-NEDD4-OCT4 pathway plays important functions during the exit of 2C-like state.

### 2C::tdTomato/OCT4-GFP reporter system can serve as a tool to uncover regulators of 2C-like state

Next, we wanted to extend the application of 2C::tdTomato/OCT4-GFP double reporter ESCs to study the regulatory mechanism underlying 2CLPT. The hypothesis is that if any factors regulate the exiting process, manipulating their expression may cause the change of Q3/Q4 ratio. We first tested whether Dux induction will change the exiting rate of 2C-like state (**Figure S6A**). Transient Dux induction at different dosage caused significant increase in the fraction of 2C-like cells with little impact on the ratio of Q3/Q4 (**Figures S6B and S6C**), suggesting that transient induction of Dux promotes the entry of 2C-like state but has little effect on the exit of 2C-like state. Indeed, the exiting rate of Q3 and Q4 was largely unchanged with different dosages of Dux induction, while the entry rate of Q2 to Q3 was significantly increased at higher dosage of Dux induction **(Figures S6D and S6E)**. We then tested another reagent, hydrogen peroxide, which previously has been shown to promote the generation of 2C-like cells (Zhang et al., 2019). When treated with hydrogen peroxide, the ratio of Q3/Q4 is significantly increased (**Figures S7A and S7B**), suggesting a delay in the exit of Q3. Indeed, we found that hydrogen peroxide delayed the exit of Q3 (**Figures S7C and S7D**). In addition, hydrogen peroxide also promoted the entry of Q2 to Q3, blocked the reversion of Q2 back to Q1, and slightly blocked the exit of Q4 to Q1 (**Figures S7C and S7D**). Together, these data show that transient Dux induction promotes the pluripotent to 2C-like state transition with little impact on the exit of 2C-like state, while hydrogen peroxide simultaneously promotes the entry of 2C-like state and blocks the exit of 2C-like state with major effect on the 2C-like to MERLOT state transition.

### Chaf1a has dual functions in regulating the entry and exit of 2C-like state

The above results support the application of 2C::tdTomato/OCT4-GFP double reporter ESCs in uncovering factors regulating the entry and exit of 2C-like state. We then performed a siRNA screening (**Figures 7A** **and S7E**) against thirteen previously identified inhibitors of 2C-like state (Fu et al., 2019; Ishiuchi et al., 2015; Li et al., 2017; Maksakova et al., 2013; Rodriguez-Terrones et al., 2017; Wu et al., 2020; Yan et al., 2019; Yang et al., 2015; Yang et al., 2020). We also included Upf1 and Smg7 as they have been previously shown to promote the exit of 2C-like state through the degradation of Dux mRNA (Fu et al., 2020). Consistently, knocking down Upf1 and Smg7 significantly increased the ratio of Q3/Q4 (**Figures 7B and 7C**), suggesting a delay in the exit of Q3. In contrast, knocking down Chaf1a significantly decreased the ratio of Q3/Q4 (**Figures 7B and 7C**), indicating that Chaf1a may promote Q3 to Q4 transition or delay Q4 to Q1 transition. In addition, knocking down other twelve genes had little impact on the ratio of Q3/Q4 (**Figures 7B and 7C**). We then sorted out Q3 and Q4 and traced the exiting process after siRNA treatment. The results showed that knocking down Chaf1a inhibits the Q4 to Q1 transition, while knocking down Upf1 or Smg7 inhibits both Q3 to Q4 and Q4 to Q1 transition (**Figures 7D and 7E**). Together with previous findings by Ishiuchi et al. (Ishiuchi et al., 2015), these data show that Chaf1a has dual functions in blocking the entry and promoting the exit of 2C-like state.

**Figure 7:**
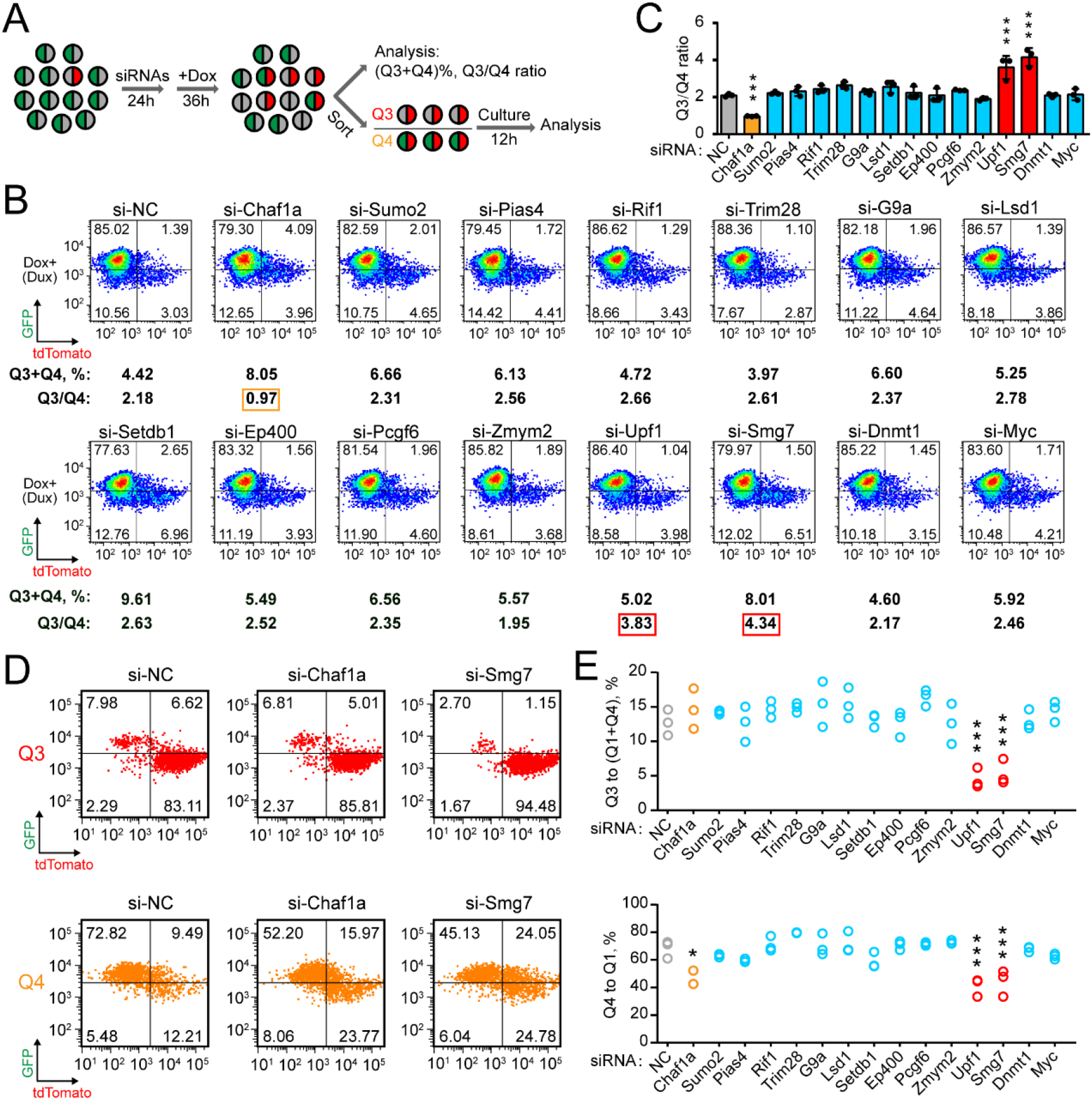
Chaf1a and NMD factors Smg7 and Upf1 promote 2CLPT at different steps. (A) Experimental design for (B)-(E). 10 nM doxycycline was used to induce Dux expression. (B and C) Flow cytometry analysis of 2C::tdTomato/OCT4-GFP reporter ESCs treated with siRNAs against candidate genes. Representative dot plots (B) with the percentage of Q3+Q4 and Q3/Q4 ratio indicated below; Quantification of Q3/Q4 ratio (C). Shown are mean ± SD, n = 3 independent experiments. The *p*-value was calculated by one-way ANOVA followed by two-tailed Dunnett’s test. (D and E) Flow cytometry analysis of dynamic transition of sorted Q3 and Q4 pre-treated with siRNAs. Representative dot plots (D); Quantification of the percentage of indicated populations (E). Each circle represents one independent experiment, n = 3. The *p*-value was calculated by one-way ANOVA followed by two-tailed Dunnett’s test.

### ATAC-Seq footprinting uncovers Klf3 as a novel transcription factor required for efficient 2CLPT

To identify more regulators during 2CLPT, we performed ATAC-Seq footprinting with TOBIAS (Bentsen et al., 2020) to screen for transcription factors with differential activity in Q3 versus Q4. As expected, Dux activity was significantly enriched in Q3 (**Figure 8A**). Interestingly, we identified multiple transcription factors from Krüpple-like family whose activities were significantly upregulated in Q4 (**Figure 8A**). We recently reported that Klf3 promotes an 8-cell like transcriptional state in mouse ESCs when overexpressed (Hao et al., 2020). Since Q4 is transcriptionally clustered with 8 and 16-cell stage embryos, we tested whether Klf3 plays a functional role in the exit of 2C-like state. Interestingly, knocking down Klf3 increased the percentage of Q3 cells and the ratio of Q3/Q4 (**Figures S8A and S8B**), suggesting a role of Klf3 in 2CLPT. Indeed, analysis of changes in sorted Q3 and Q4 cells showed that knocking down Klf3 significantly delayed the exit of 2C-like state at both 2C-like-to-MERLOT and MERLOT-to-pluripotent state transition steps (**Figures 8B and 8C**). In contrast, knocking down Klf4, a known pluripotency transcription factor, had little impact on the exit of 2C-like state (**Figures S8C and S8D**). Furthermore, transfection of synthetic BFP-t2a-Klf3 mRNA significantly promoted the exit of 2C-like state (**Figure 8D**). We noticed that BFP-t2a-Klf3 mRNA converted more Q1 cells to Q2 (**Figures S8E and S8F**), indicating the importance of temporal control of KLF3 expression or activity during the exit of 2C-like state. Together, these data identifies Klf3 as a novel transcription factor required for efficient 2CLPT.

**Figure 8:**
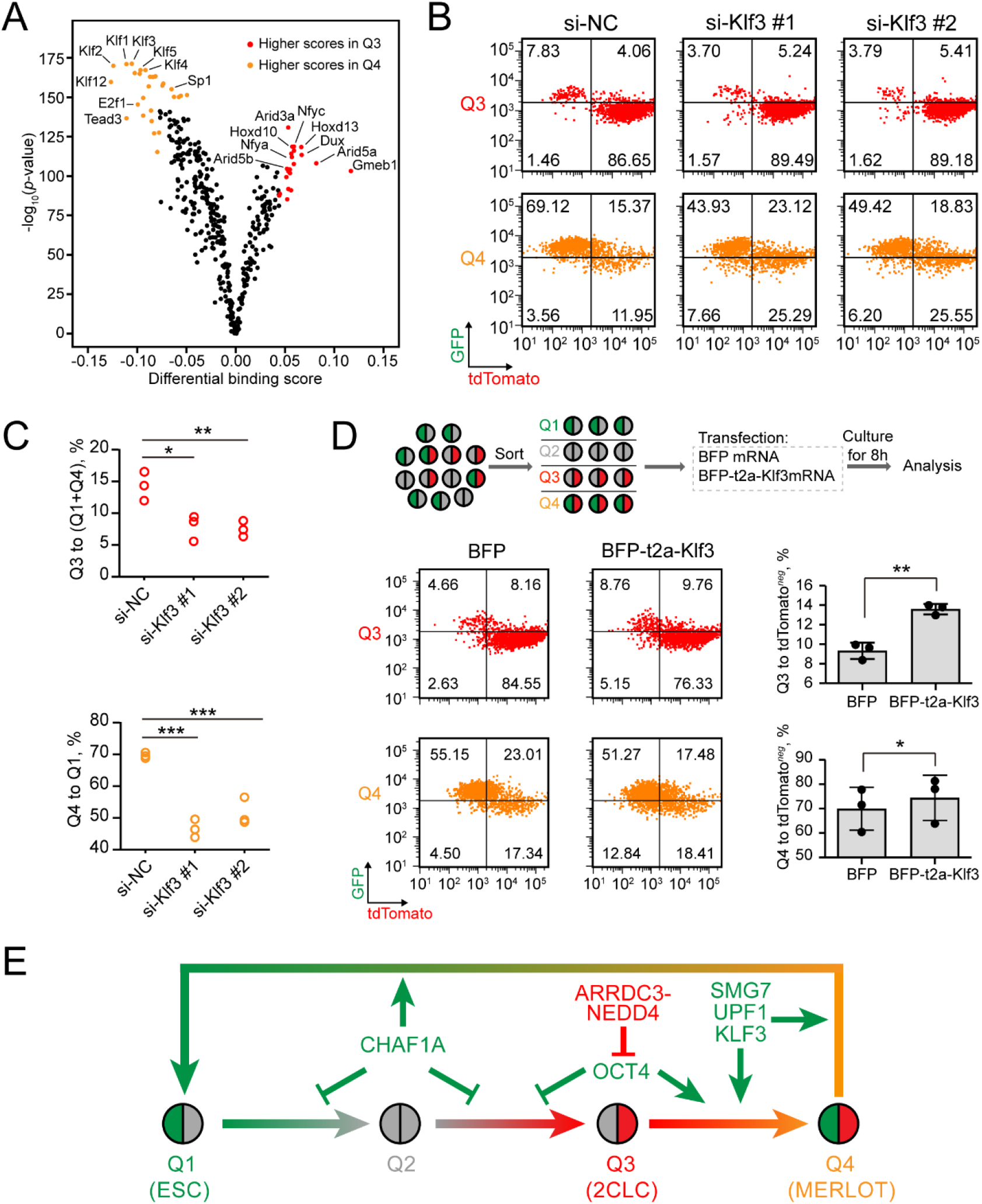
ATAC footprinting identifies Klf3 as a novel transcription factor required for efficient 2CLPT. (A) Volcano plot showing differential binding activity of all investigated transcription factors (TFs) between Q3 and Q4. Each dot represents one TF motif. (B and C) Flow cytometry analysis of dynamic transition of sorted Q3 and Q4 pre-treated with control and Klf3 siRNAs. Representative dot plot (B); Quantification of the percentage of indicated populations (C). Each circle represents one independent experiment, n = 3. The *p*-value was calculated by one-way ANOVA followed by two-tailed Dunnett’s test. (D) Flow cytometry analysis of dynamic transition of sorted Q3 and Q4 transfected with BFP and BFP-t2a-Klf3 mRNAs. Upper, experimental design; Lower left, representative dot plot; Lower right, quantification of the percentage of indicated populations. Each circle represents one independent experiment, n = 3. The *p*-value was calculated by paired two-tailed Student’s *t* test. (E) Summary graph showing multiple regulators controlling the entry and exit of 2C-like state at different steps.

## Discussion

The molecular mechanism underlying the exit of 2C-like state is largely unclear. Identification of an intermediate state that can be labeled and isolated through genetically encoded reporters will provide a powerful tool to study the exit of 2C-like state. Here we identify an intermediate state during the exit of 2C-like state that can be labeled by the co-expression of 2C::tdTomato and OCT4-GFP. Unlike conventional reporters only reflecting the promoter activity, OCT4-GFP reporter integrates the input from the regulation of promoter activity, mRNA stability and protein stability. Since post-transcriptional and post-translational regulation are widespread during cell fate change (Chua et al., 2020; Lu et al., 2009), the design strategy of our OCT4-GFP fusion protein reporter may have implications in uncovering and isolating rare cell types in other biological settings such as stem cell differentiation and cancer.

Our study shows that 2CLPT involves a two-step process with the first step being 2C-like-to-MERLOT state transition and the second step being MERLOT-to-pluripotent state transition. The MERLOT state is likely different from an intermediate state that is identified through transcriptomic analysis by Fu et al. (Fu et al., 2020), since cells in their intermediate state are negative for 2C::tdTomato. Therefore, 2CLPT could be a multiple-step process that involves two or more intermediate states. The verification of this hypothesis requires the development of a genetically encoded reporter for the intermediate state identified by Fu et al. (Fu et al., 2020). More interestingly, the MERLOT state clustered closely with 8 and 16-cell stage embryos at both transcriptome and epigenome levels, suggesting that 2CLPT may mimic some of molecular features during early mouse embryo development. Therefore, investigation of 2CLPT could help identify potential regulators of early embryo development.

Oct4 is a transcription factor that plays important roles in the maintenance of pluripotent state and reprogramming (Jaenisch and Young, 2008; Nichols et al., 1998; Radzisheuskaya and Silva, 2014; Shi and Jin, 2010; Takahashi and Yamanaka, 2006). We found that OCT4 protein is significantly repressed by ARRDC3-NEDD4-proteasome degradation pathway during the entry of 2C-like state. In addition, ARRDC3 knockout or OCT4 overexpression significantly accelerated the 2C-like-to-MERLOT state transition to a similar degree. However, ARRDC3 knockout or ARRDC3/4 double knockout did not completely eliminate OCT4-GFP^-^ cells, suggesting other redundant or parallel mechanisms repressing OCT4 expression. In addition, we confirmed Fu et al.’s finding that NMD factors Smg7 and Upf1 promote the exit of 2C-like state (Fu et al., 2020). Nevertheless, by the merit of our reporter system, we provided a more comprehensive understanding on the role of Smg7 and Upf1 in both 2C-like-to-MERLOT and MERLOT-to-pluripotent state transition steps. In addition, through re-evaluating previously identified inhibitors of 2C-like state and footprinting analysis of ATAC-Seq, we discovered Chaf1a and Klf3 as two important regulators required for efficient 2CLPT. Interestingly, Kinisu et al. find that the binding motif of Klf3 is enriched at LTR elements of MERVL, ORR1A0 and ORR1A1 (Kinisu et al., 2021). Therefore, Klf3 may directly repress the expression of ERVs by recruiting a cohort of other corepressors (Pearson et al., 2011; Turner and Crossley, 1998). In addition, knockdown of histone chaperone Chaf1a delayed the MERLOT-to-pluripotent state transition but not the 2C-like-to-MERLOT state transition. Therefore, our results and results from Ishiuchi et al. (Ishiuchi et al., 2015) together support a model that proper nucleosome assembly during replication safeguards ESC identity by blocking the emergence of 2C-like cells and reinstates the ESC fate at later stage of 2CLPT. Altogether, these data demonstrate that our double reporter system is a powerful tool to identify regulators of 2CLPT. Mechanistic and functional dissection of these regulators could provide more insights on the early embryo development around ZGA.

We propose to identify regulators of 2CLPT based on the alteration of Q3/Q4 ratio. However, if a treatment similarly affects 2C-like-to-MERLOT and MERLOT-to-pluripotent state transition, the change of Q3/Q4 ratio may not be observed. In these rare scenarios, dynamic cell fate changes of sorted 2C-like and MERLOT cells after treatments need be carefully analyzed before any conclusions can be made.

In conclusion, we have identified a genetically isolatable intermediate state during the exit of 2C-like state. Furthermore, we have confirmed Smg7 and Upf1 and uncovered ARRDC3-NEDD4-OCT4 pathway, Chaf1a and Klf3 as essential regulators of 2CLPT at different steps (**Figure 8E**). Our study provides a valuable tool to study 2CLPT and have implications for isolating rare cell types in other biological processes.

## Materials and Methods

### Cell culture

Mouse ESCs were cultured in Serum/LIF medium composed of KnockOut^TM^ DMEM (Gibco, 10829081) supplemented with 15% FBS (Hyclone, H3007103), 1 mM L-glutamine (Gibco, 25030081), 0.1 mM non-essential amino acids (NEAA) (Gibco, 11140050), 1 X Penicillin/Streptomycin (GIBCO, 15140163), 0.1 mM 2-mercaptoethanol and 1,000 U/ml mouse leukemia inhibitory factor (mLIF). Cells were passaged every other day and the medium were refreshed daily. Mycoplasma nucleic acids were routinely tested by PCR to ensure mycoplasma free culture conditions. For hydrogen peroxide treatment assay, 50 µM, 100 µM, 200 µM hydrogen peroxide was freshly prepared in ES medium without 2-mercaptoethanol. After 48 h treatment, cells were collected for cytometry analysis. For kinetics analysis assay, Q1/2/3/4 were first sorted out, then treated with freshly prepared 100 µM or 200 µM hydrogen peroxide in ES medium without 2-mercaptoethanol. After 12 h culture, Q1/2/3/4 were collected for flow cytometry analysis.

### Construction of MERLOT reporter cell line

2C::tdTomato reporter ESC line was constructed as previously described (Yan et al., 2019). OCT4-EGFP reporter was constructed in 2C::tdTomato cell line. sgRNA (5’-TCTCCCATGCATTCAAACTG-3’) was designed to direct Cas9 to make a double-strand break near the stop codon of Oct4. A donor plasmid containing EGFP sequence flanked by 1,000 bp homology arms was co-transfected with Cas9-sgRNA. ESCs were selected in 300 μg/ml Hygromycin B for 3 days. EGFP positive cells were sorted by FACS and seeded at clonal density. Single knock-in colonies were screened by genotyping and expanded for further experiments. For Dux overexpression, the expression plasmid containing Dux under the control of TRE3G promoter was transfected into MERLOT reporter cell line through PiggyBac transposon system.

To induce the transition of ESCs to 2C-like cells, MERLOT reporter ESCs were treated with doxycycline for 2 h to induce Dux expression at concentrations indicated. After withdrawal of doxycycline, MERLOT reporter ESCs were cultured for additional 24 h or 36 h before analysis or sorting. For siRNA knockdown assays, siRNAs were transfected 1 day before the treatment of doxycycline.

### RNA extraction and RT-qPCR

Total RNA was extracted using Trizol reagent following standard procedures. cDNAs were obtained by reverse transcription from 500 ng total RNA using HiScript II Q RT SuperMix for qPCR (Vazyme, R223). Quantitative PCR (qPCR) was carried out using AceQ qPCR SYBR Green Mater Mix reagent (Vazyme, Q141) on StepOne Plus Real-Time PCR System (Applied Biosystems). Quantification was performed through comparative cycle threshold (CT) method normalized to β-actin.

### CRISPR knock out and CRISPRa

To knock out Arrdc3 and Arrdc4, paired sgRNAs were designed by https://www.benchling.com/crispr/ to target Arrdc3 exon2 and Arrdc4 exon1 to make frame shift. The paired sgRNAs for Arrdc3 were 5’-CCACAGACGTAAGTATTCTA-3’ and 5’-GCCTTCCTGAATGAATAGTG-3’, for Arrdc4 were 5’-CTACTCAAGCGGCGAGACAG-3’ and 5’-CCTGCGGTTGAGTCTGCTGG-3’. Knockout was verified by genomic PCR followed by Sanger sequencing and RT-qPCR. For CRISPRa screening, ubiquitin-proteasome pathway related genes were picked based on information from https://ubihub.thesgc.org/static/UbiHub.html, as well as gene annotation from https://www.uniprot.org/. sgRNAs targeting candidate genes were designed by http://crispr-era.stanford.edu/. In principle, sgRNAs for CRISPRa were designed to target the -200 to 0 bp upstream of transcription start site.

### siRNA design and transfection

siRNAs in this study were designed by https://horizondiscovery.com/en/ordering-and-calculation-tools/sidesign-center or collected from references. JetPRIME® (Polyplus Transfection, PT-114-75) was used for siRNA transfection according to the manufacturer’s instruction. Final concentration of siRNAs was 50 nM.

For siRNA treatment experiments in Figures 7A-E, 8B, 8C and S8A-D, fresh medium was changed 10 h after siRNA transfection. 24 h after transfection, doxycycline was added at final concentration of 10 nM transiently for 2 h to induce Dux expression. 36 h after Dux induction, cells were collected for flow cytometry analysis or cell sorting. For transition kinetics analysis, the sorted Q1/2/3/4 were cultured for 12 h before flow cytometry analysis. For siRNA treatment experiments in Figures S5F-H, siRNAs were transfected 48 h before flow cytometry analysis. 2.5 µM doxycycline was added throughout the experiments from cell seeding to siRNA treatment to induce Arrdc3 expression.

### In vitro transcription and transfection of mRNA

mRNAs were synthesized by in vitro transcription (IVT) using HiScribe^TM^ T7 ARCA mRNA Kit (NEB, E2060) according to the manufacturer’s instruction with slight modification. In brief, the complementary DNA (cDNA) encoding desired mRNA was cloned under the control of T7 promoter. The plasmid was linearized by PmeI or XbaI restriction enzyme. After purification by phenol-chloroform extraction and ethanol precipitation, the linearized plasmids were used as templates for in vitro transcription by T7 polymerase. Synthesized mRNAs were purified by lithium chloride precipitation and dissolved in nuclease-free water. Lipofectamine MessengerMAX™ (Invitrogen, LMRNA001) was used for mRNA transfection. After seeding sorted cells into 24-well plate, mRNA transfection was immediately conducted for cells in suspension. 250 ng mRNA with 0.5 µl MessengerMAX reagent were used per well.

### Flow cytometry and cell sorting

Cells were digested into single cells by 0.1% trypsin in 37℃, 10 min and resuspended in ice-cold DMEM medium. Cell suspensions were filtered through 40µm strainer and then analyzed on BD LSRFortessa SORP. Cell sorting was performed on BD FACSAria III or BD FACSAria SORP. Data analysis was performed using FlowJo software.

### Live-cell imaging

Live-cell imaging was carried out on a Dragonfly spinning disk confocal microscope (Andor Technologies) equipped with Okolab boldline incubator (Okolab) and mounted on a Leica DMI8 using 40 X objective (Leica Camera). Focus stability was maintained using the Perfect Focus System (PFS) during imaging. Cells were imaged using 488 nm laser line and 562 nm laser line both at 200 ms exposure and 5% transmission, along with DIC to capture bright-field image. To trace exit process of 2C-like cells, Q3 cells were sorted out by FACS and plated on 35 mm glass-bottom dish (Cellvis, D35-20-1-N) coated with 0.2% gelatin 8 h before imaging. Images were collected at a frame rate of 15 min with a duration of 14 h. Data were processed and analyzed using Fiji software.

### Immunostaining

Cells were fixed with 4% PFA at room temperature for 20 min, then permeabilized with 0.25% Triton X-100 for 15 min and blocked with 3% BSA, 1% FBS in PBS at 37℃ for 1 h. Primary antibodies were diluted into blocking buffer at recommended concentrations and incubated with cells at 37℃ for 2 h. After three times washing with PBS, the cells were incubated with Alexa Fluor secondary antibody at room temperature for 1 h. Finally, cells were stained with DAPI before imaging on microscope. For flow cytometry analysis, cells were first digested into single cells and then fixed with 4% PFA at room temperature for 20 min, followed by incubation in ice-cold 90% methanol at 4 ℃ for 15 min. The procedures of blocking, primary/secondary antibodies incubation and DAPI staining were similar to that of attached cells. Flow cytometry analysis was performed on BD LSRFortessa SORP device.

### Western blot

Cells were lysed on ice for 30 min using Triton X-100 lysis buffer containing 20 mM Tris-HCl (pH 7.4), 150 mM NaCl, 1 mM EGTA, 1 mM Na_2_EDTA, 2.5 mM Na_4_P_2_O_7_, 1 mM β-glycerophosphate and 1% Triton X-100 supplemented with EDTA-free protease inhibitors (Thermo, A32963). Cell lysates were centrifuged at 12,000 rpm for 30 min at 4 ℃. SDS-PAGE electrophoresis was conducted following standard procedures. Primary antibodies used in this study were mouse anti-OCT4 (Santa Cruz, sc-5279), rabbit anti-OCT4 (CST, 83932), anti-SOX2 (Millipore, ab5603), anti-FLAG (Sigma, F1804), anti-GAPDH (Proteintech, 60004), anti-HA (CST, C29F4). Secondary antibodies were IRDye 800CW donkey-anti-rabbit (LI-COR, 926-32213), IRDye 680LT donkey-anti-mouse (LI-COR, 926-68022), IRDye 800CW goat-anti-mouse (LI-COR, 926-32210). PVDF membranes were imaged using Odyssey colour infrared laser scan-imaging instrument and obtained images were processed and analyzed by ImageJ.

### Immunoprecipitation and ubiquitination detection

Cells were lysed on ice for 30 min using Triton X-100 lysis buffer and centrifuged at 12,000 rpm for 30 min at 4 ℃. Primary antibodies were first incubated with protein G Dynabeads (Invitrogen, 10003) at room temperature for 10 min on a rotator. Primary antibodies used in this study were mouse anti-OCT4 (Santa Cruz, sc-5279), rabbit anti-OCT4 (CST, 83932), anti-FLAG (Sigma, F1804), anti-HA (CST, C29F4). For immunoprecipitation, cell lysates were incubated with protein G at 4 ℃ for 12-16 h on a nutator. Immunoprecipitates were then washed three times with Triton X-100 lysis buffer. Whole cell lysates and immunoprecipitates were analyzed by western blot. For ubiquitination detection, HA-Ub expression construct was stably transfected and MG132 was added in culture medium 4 h before lysis.

### RNA-Seq library preparation and sequencing

RNA-seq was performed using VAHTSTM mRNA-seq V3 Library Prep Kit (Vazyme, NR611) according to the manufacturer’s instruction. In brief, poly(A)^+^ mRNAs were captured by poly-Oligo dT magnet beads followed by fragmentation by bivalent cation buffer at 94 ℃ for 5min. ds-cDNAs were then synthesized and ligated to adaptors. After PCR amplification, the library was subjected to sequence on an Illumina Nova-seq 6000 sequencing system.

### ATAC-Seq library preparation and sequencing

ATAC-Seq was performed as previously described with slight modifications (Buenrostro et al., 2013). 50,000 cells were sorted out by FACS and collected by centrifugation at 500 g for 5 min at 4 ℃. After washing with PBS, cell pellet was resuspended in 50 μl of cold lysis buffer (10 mM Tris-HCl, pH 7.4, 10 mM NaCl, 3 mM MgCl2, 0.1% IGEPAL CA-630) for 5 min at 4 ℃. Centrifuge immediately after lysis at 500 g for 10 min at 4 ℃ and the cell pellets were resuspended with 50 μl transposition mix (10 μl 5 X TTBL, 5 μl TTE Mix V50, 35 μl nuclease-free H_2_O, Vazyme TD501). Mixture was incubated at 37 ℃ for 30 min followed by purification with a Zymo DNA Clean and Concentrator-5 Kit (Zymo, DP4033). Transposed DNA was eluted in 24 μl elution buffer and amplified by PCR reaction (10 μl 5 X TAB, 5 μl PPM, 5 μl N5 Primer, 5 μl N7 Primer, 1 μl TAE, Vazyme TD501) for 13 cycles. Amplification products were further purified by 1.6 X VAHTS DNA Clean Beads (Vazyme, N411) with 0.6 X-1 X dual size selection. The libraries were then subjected to sequence by an Illumina Nova-seq 6000 sequencing system.

### CUT&Tag library preparation and sequencing

CUT&Tag was performed as previously described with slight modifications (Kaya-Okur et al., 2019). 100,000 cells were sorted out by FACS and resuspended in 100 μl 1 X wash buffer (20 mM HEPES pH 7.5, 300 mM NaCl, 0.5 mM spermidine with EDTA-free protease inhibitors). With gentle vortex, 10 μl ConA beads slurry were added dropwise and incubated at room temperature for 10 min. After collection by magnet stand, primary antibodies (1:100 or as recommended) in 100 μl ice-cold antibody buffer (20 mM HEPES pH 7.5, 300 mM NaCl, 0.5 mM spermidine, 0.025% digitonin, 2mM EDTA, 0.1% BSA with EDTA-free protease inhibitors) were added: H3K27ac (Abcam, ab4729), H3K9me3 (Abcam, ab8898), H2AK119ub (CST, 8240), rabbit IgG (Sigma, I8140), and incubated at room temperature for 2 h on a nutator. Then incubated with secondary antibody (Abcam, ab6702, 1:100) in Dig-wash buffer (20 mM HEPES pH 7.5, 300 mM NaCl, 0.5 mM spermidine, 0.025% digitonin with EDTA-free protease inhibitors) at room temperature for 1 h. 100 μl 0.04 μM hyperative pG-Tn5 (Vazyme, TD901) in Dig-300 buffer was added and incubated at room temperature for 1 h. Tagmentation was carried out by adding 10 mM MgCl_2_ and incubation at 37 ℃ for 1 h, then stopped by adding 16.6 mM EDTA, 0.1% SDS and 16.6 μg/mL proteinase K and incubation at 50 ℃ for 1 h. Tagmentated DNA was extracted by phenol chloroform extracting method and dissolved in 24 μl nuclease-free H_2_O, then amplified by PCR reaction (10 μl 5 X TAB, 5 μl PPM, 5 μl P5 Primer, 5 μl P7 Primer, 1 μl TAE, 5 μl H_2_O, Vazyme TD901) for 16 cycles. Amplification products were further purified by 1.2 X VAHTS DNA Clean Beads (Vazyme, N411) and subjected to sequence by an Illumina Nova-seq 6000 sequencing system.

### RNA-Seq data processing and analysis

Sequencing reads were processed by Trim Galore to trim low quality reads and remove adapters. Processed reads were then aligned to the mouse genome (mm10) with STAR (version 2.5.0) using the GENCODE transcript annotation as transcriptome guide. All programs were processed following default settings except for special annotation. The TPM values generated by StringTie (version 1.1.2) were used to quantify the expression level. Differentially expressed genes were determined by DESeq2. The enrichment of selected gene sets was calculated by java GSEA Desktop Application. GO enrichment analysis was performed by DAVID (v6.8). R software (v3.5.1) was used for the generation of scatter plot, box plot, violin plot, heatmap and PCA analysis. For the comparison with the pre-implantation mouse embryo, data were downloaded from GEO (GSE45719) (Deng et al., 2014). For the comparison with the intermediates identified by Fu et al., data were downloaded from GEO (GSE133234 and GSE121459) (Fu et al., 2020; Fu et al., 2019). Batch-effect was removed by R package QuantNorm.

### ATAC-Seq data processing and analysis

Sequencing reads were processed by Trim Galore to trim low quality reads and remove adapters. Processed reads were then aligned to the mouse genome (mm10) with STAR (version 2.5.0). ATAC-Seq peaks were called using MACS2 ‘callpeak’ (-B –nomodel --nolambda --shift −100 --extsize 200). For the comparison with ATAC profile in the pre-implantation mouse embryo, data were downloaded from GEO (GSE66390) (Wu et al., 2016) and re-processed as described above. MACS2 ‘bdgdiff’ subcommand (−l 300 –g 250) was used to call “differential peaks” between 4C and 8C embryo. The deepTools was used to plot heatmaps and metagene profiles. Footprinting signatures of transcription factors were calculated by TOBIAS (Bentsen et al., 2020).

### CUT&Tag data processing and analysis

Sequencing reads were processed by CUT&Tag data processing and analysis pipeline (https://yezhengstat.github.io/CUTTag_tutorial/). Igvtools was used to visualize the high-throughput sequencing signals.

### Quantification and statistical analysis

No statistical methods were used to pre-determine the sample size. No data were excluded from the analyses. Statistical analyses were performed by GraphPad Prism 8 and R software version 4.0.3. Details of statistics were listed in corresponding figure legends. Significance defined as *p*-value was indicated on figures (* *p*-value < 0.05, ** *p*-value < 0.01, *** *p*-value < 0.001, **** *p*-value < 0.0001).

### Data and code availability

RNA-Seq, ATAC-Seq and CUT&Tag data that support the findings of this study will be deposited in the Gene Expression Omnibus (GEO) database. Software/packages used to analyze the dataset are either freely or commercially available. All other data and code supporting the findings of this study are available from the corresponding author upon reasonable request.

## Acknowledgments

We would like to thank members of Wang laboratory and Dr. Wei Xie for critical reading and discussion of the manuscript. We thank Huan Yang and Yinghua Guo at National Center for Protein Sciences, Peking University for assistance with cell sorting. We thank Fei Wang and Xin Li at the Center for Quantitative Biology, Peking University for assistance with flow cytometry analysis and live cell imaging, respectively. We thank Dr. Siying Qin at the Core Facilities of the School of Life Sciences, Peking University for assistance with optical/confocal imaging. This study was supported by The National Key Research and Development Program of China [2018YFA0107601] and the National Natural Science Foundation of China [91940302 and 32025007] to YW.

## Author contributions

CZ and MS performed all experiments with help from others. JH performed all bioinformatics analysis. YXL helped with constructing plasmids and cell culture experiments. All authors were involved in the interpretation of data. YW conceived and supervised the project. YW and CZ wrote the manuscript with help from JH, MS and WY.

---

**Figure S1:**
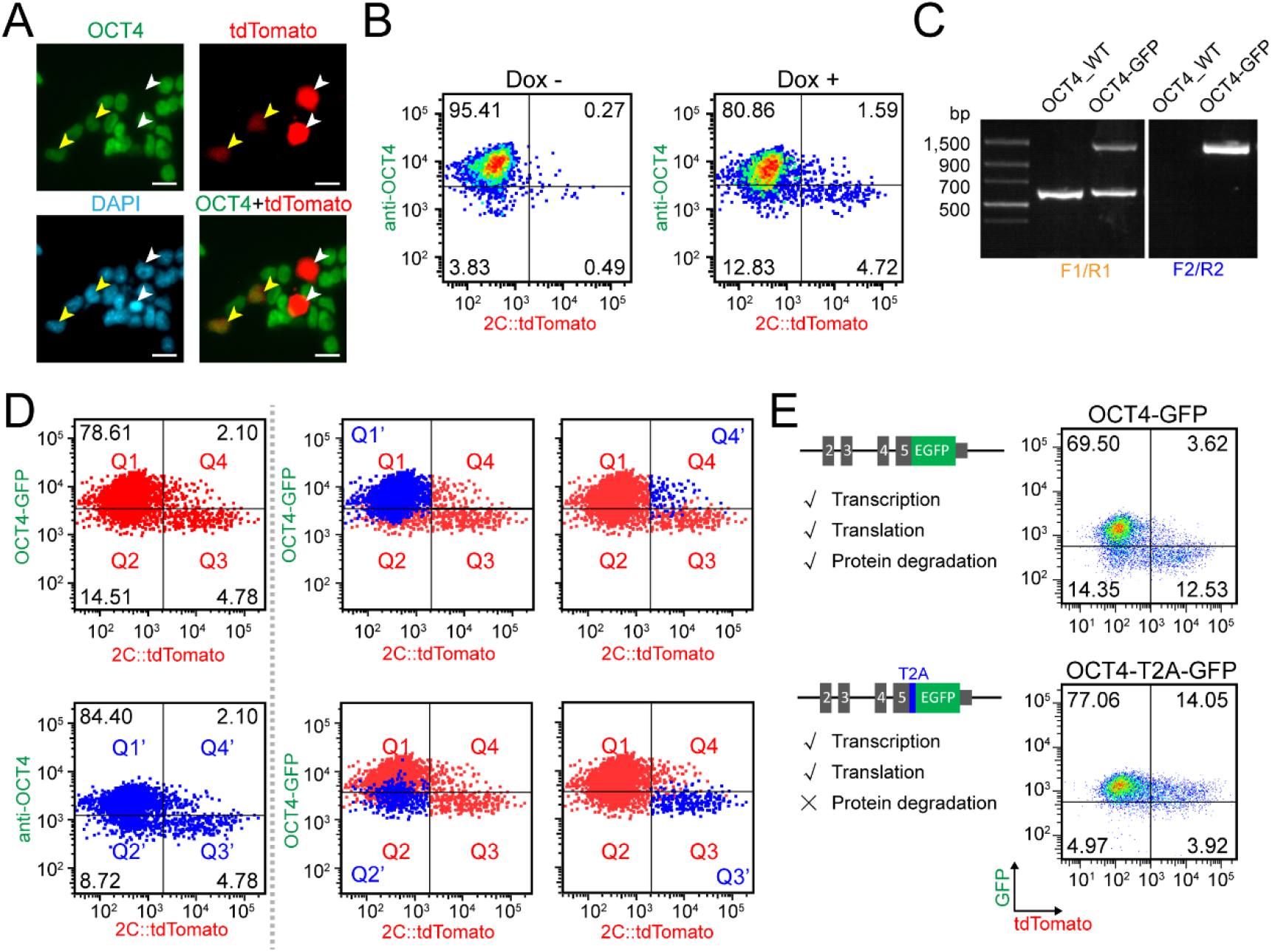
Identification of 2C::tdTomato^+^OCT4-GFP^+^ cells in mouse ESCs. Related to Figure 1. (A) Immunostaining of OCT4 protein in 2C::tdTomato reporter ESCs. White arrows point to 2C::tdTomato^+^OCT4^-^ cells; Yellow arrows point to 2C::tdTomato^+^OCT4^+^ cells. Scale bars, 20 µm. (B) Flow cytometry analysis of 2C::tdTomato reporter ESCs immunostained by OCT4 antibody. 20 nM doxycycline was used to induce Dux expression in the Dox+ sample. (C) PCR-based genotyping to validate GFP knock-in at *Oct4* locus. Two paired primers (F1/R1, F2/R2) were designed for genotyping, which are indicated in Figure 1A. (D) Flow cytometry analysis of 2C::tdTomato/OCT4-GFP reporter ESCs immunostained by OCT4 antibody. 20 nM doxycycline was used to induce Dux expression. Left, representative dot plots; ESCs were divided into Q1/2/3/4 by *y* axis: OCT4-GFP, *x* axis: 2C::tdTomato, and into Q1’/2’/3’/4’ by *y* axis: anti-OCT4, *x* axis: 2C::tdTomato; Right, Q1’/2’/3’/4’ were projected onto Q1/2/3/4. (E) Flow cytometry analysis of 2C::tdTomato/OCT4-GFP and 2C::tdTomato/OCT4-T2A-GFP reporter ESCs. 40 nM doxycycline was used to induce Dux expression. Left, reporter design, gene regulation steps that can be monitored by each reporters are indicated; Right, representative dot plots.

**Figure S2:**
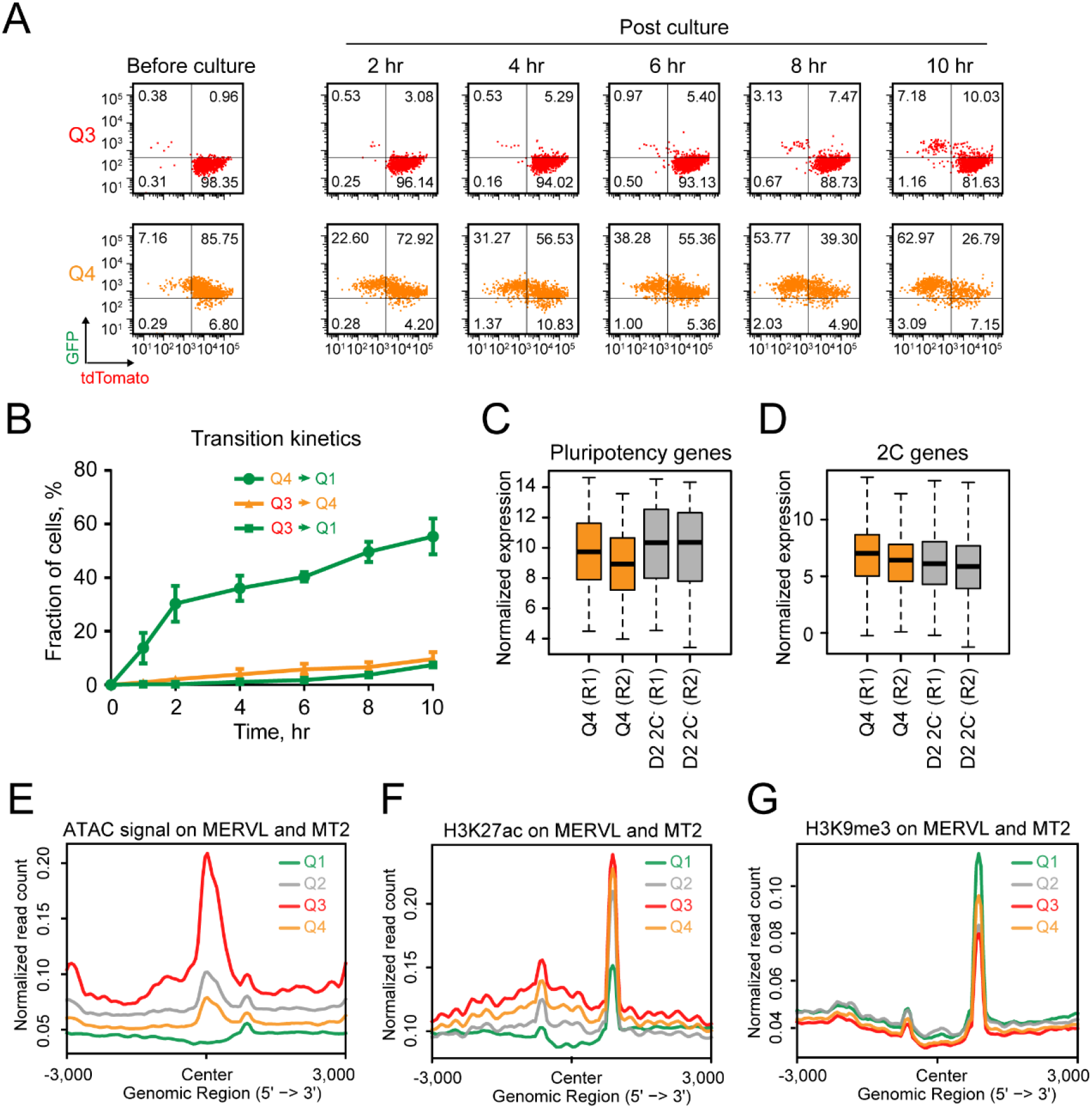
Characterization of the dynamic transition, transcriptome and epigenome of MERLOT cells. Related to Figures 1 and 3. (A and B) Flow cytometry analysis of the dynamic transition of sorted Q3 and Q4 over time. Representative dot plots (A); Quantification of the percentage of indicated populations (B). Shown are mean ± SD, n = 3 independent experiments. (C and D) Box and whisker plot showing the expression level of (C) pluripotency genes (n=43) and (D) 2C specific genes (n=333) in Q4 and D2 2C^-^. Center line, median; Box limits, upper and lower quartiles; Whiskers, 1.5× interquartile range. The RNA-Seq data of D2 2C^-^ are Fu et al.’s study (Fu et al., 2020). (E-G) Read-count tag density pileups of (E) ATAC signal, (F) H3K27ac profile and (G) H3K9me3 profile on MERVL-int and MT2_Mm.

**Figure S3:**
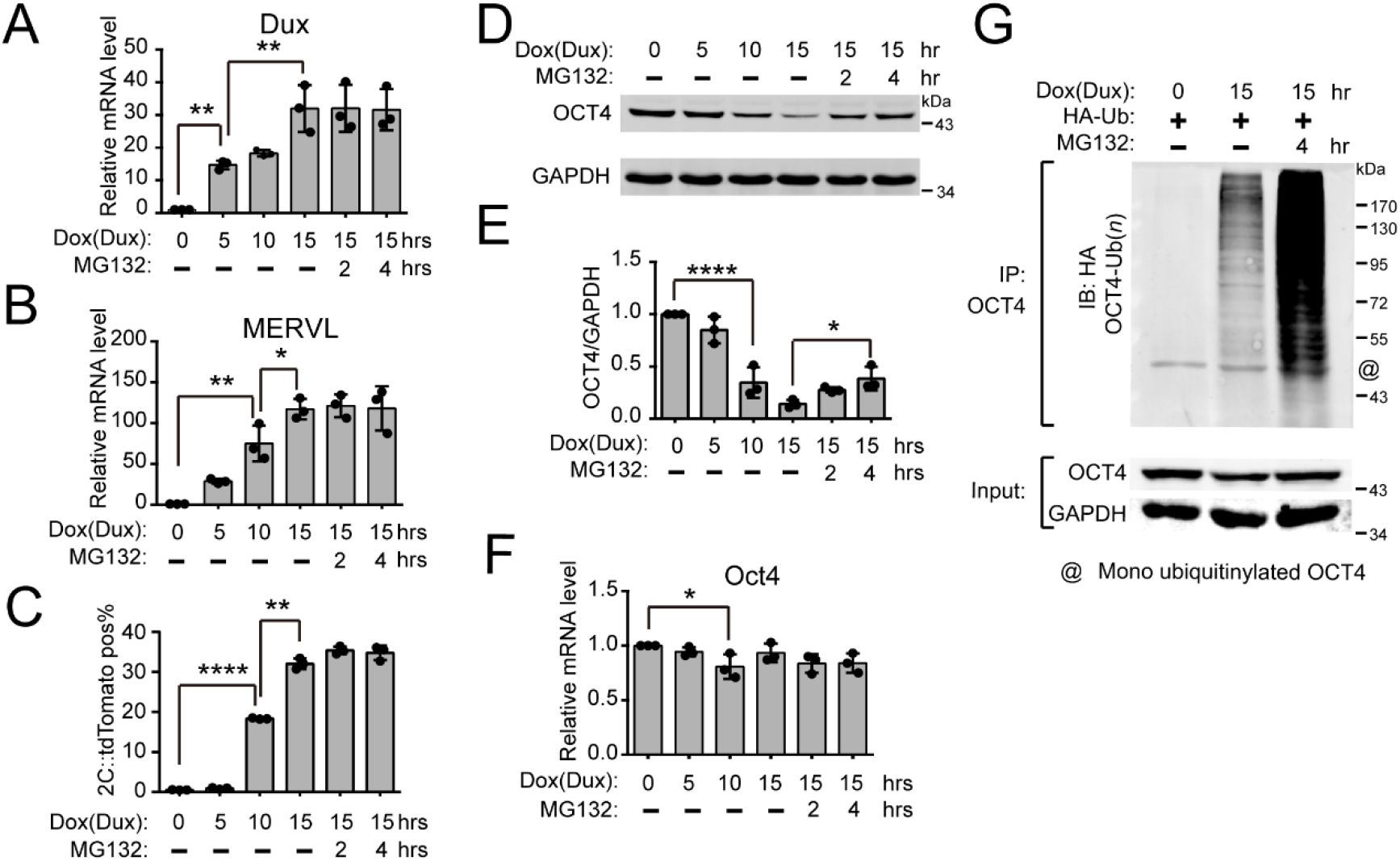
OCT4 protein is degraded through ubiquitin-proteasome pathway during the entry of 2C-like state. Related to Figure 6. (A and B) RT-qPCR analysis of the expression of (A) Dux and (B) MERVL upon Dux induction by 40 nM doxycycline over time. MG132 was used to inhibit proteasome activity. Shown are mean ± SD, n = 3 independent experiments. The *p*-value was calculated by one-way ANOVA followed by two-tailed Tukey’s test. (C) Flow cytometry analysis of the percentage of 2C::tdTomato^+^ cells upon Dux induction by 40 nM doxycycline over time. Shown are mean ± SD, n = 3 independent experiments. The *p*-value was calculated by one-way ANOVA followed by two-tailed Tukey’s test. (D-F) The expression of Oct4 mRNA and protein upon Dux induction by 40 nM doxycycline over time. Western blot analysis of OCT4 protein (D); Quantification of OCT4 protein (E). Shown are mean ± SD, n = 3 independent experiments. The *p*-value was calculated by one-way ANOVA followed by two-tailed Tukey’s test. RT-qPCR analysis of Oct4 mRNA (F); shown are mean ± SD, n = 3 independent experiments. The *p*-value was calculated by one-way ANOVA followed by two-tailed Tukey’s test. (G) Western blot analysis of OCT4 ubiquitination in OCT4 immunoprecipitated samples upon Dux induction by 40 nM doxycycline with or without MG132 treatment.

**Figure S4:**
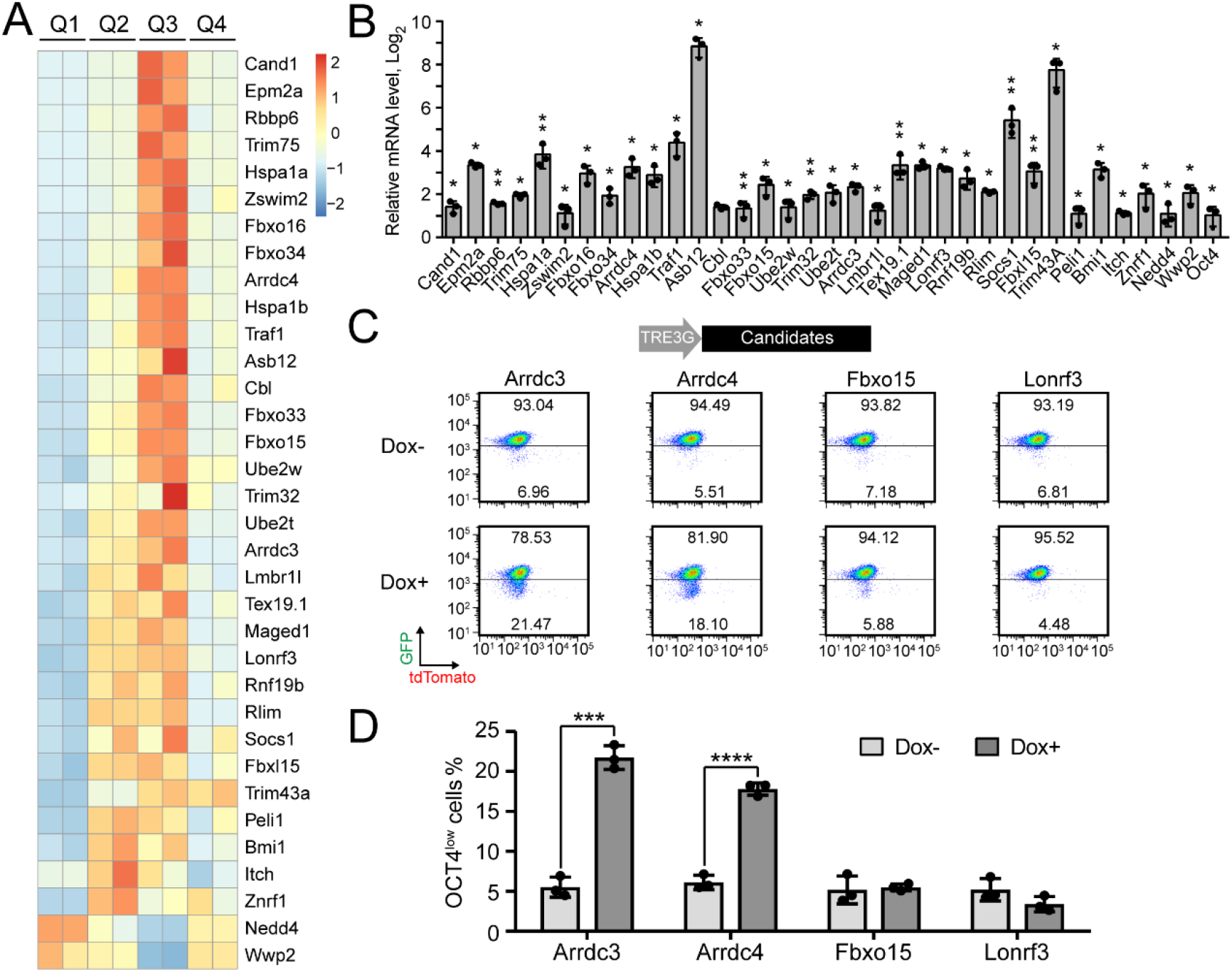
CRISPRa screen identifies Arrdc3 and Arrdc4 potentially regulating the degradation of OCT4 protein. Related to Figure 6. (A) Heatmap showing the expression of ubiquitin pathway related genes highly expressed in Q2 and Q3. (B) RT-qPCR analysis of upregulation of candidate genes by CRISPRa. Shown are mean ± SD, n = 3 independent experiments. The *p*-value was calculated by upaired one-tailed Student’s *t* test. (C and D) Flow cytometry analysis of OCT4-GFP reporter with or without Arrdc3, Arrdc4, Fbxo15, Lonrf3 induction by 2.5 µM doxycycline. Representative dot plot (C); Quantification of the percentage of OCT4-GFP*^low^* population (D). Shown are mean ± SD, n = 3 independent experiments. The *p*-value was calculated by unpaired two-tailed Student’s *t* test.

**Figure S5:**
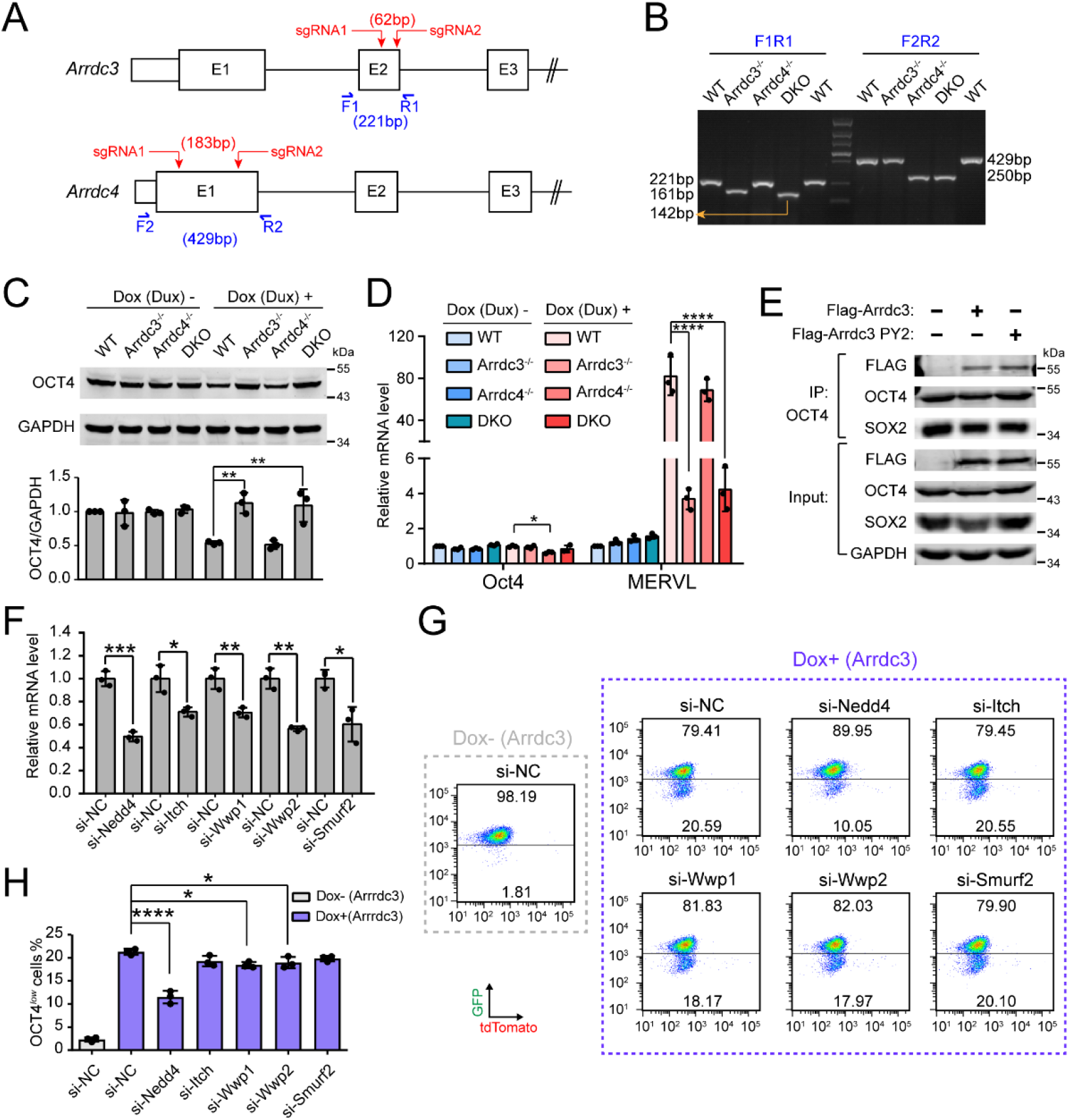
ARRDC3-NEDD4-proteasome pathway regulates OCT4 protein stability. Related to Figure 6. (A) Schematic of the knockout strategy of *Arrdc3* and *Arrdc4*. (B) PCR-based genotyping to validate the knockout of *Arrdc3* and *Arrdc4*. Two paired primers (F1/R1, F2/R2) were designed for genotyping, which are indicated in (A). (C) Western blot analysis of OCT4 protein in wild-type, *Arrdc3* knockout, *Arrdc4* knockout and *Arrdc3*/*Arrdc4* double knockout ESCs with or without Dux induction by 40 nM doxycycline. Upper, representative images of western blot; Lower, quantification of OCT4 protein. Shown are mean ± SD, n = 3 independent experiments. The *p*-value was calculated by one-way ANOVA followed by two-tailed Tukey’s test. (D) RT-qPCR analysis of the expression of Oct4 and MERVL in wild-type, *Arrdc3* knockout, *Arrdc4* knockout and *Arrdc3*/*Arrdc4* double knockout ESCs with or without Dux induction by 40 nM doxycycline. Shown are mean ± SD, n = 3 independent experiments. The *p*-value was calculated by one-way ANOVA followed by two-tailed Tukey’s test. (E) Western blot analysis of OCT4 immunoprecipitated samples with or without Arrdc3 or Arrdc3 PY2 overexpression. (F-H) Flow cytometry analysis of OCT4-GFP reporter ESCs treated with siRNAs against Nedd4, Itch, Wwp1, Wwp2 and Smurf2. 2.5 µM doxycycline was used to induce Arrdc3. RT-qPCR analysis of the efficiency of siRNAs (F). Shown are mean ± SD, n = 3 independent experiments. The *p*-value was calculated by unpaired two-tailed Student’s *t* test. Representative dot plot (G); Quantification of the percentage of OCT4-GFP*^low^* population (H). Shown are mean ± SD, n = 3 independent experiments. The *p*-value was calculated by one-way ANOVA followed by two-tailed Dunnett’s test.

**Figure S6:**
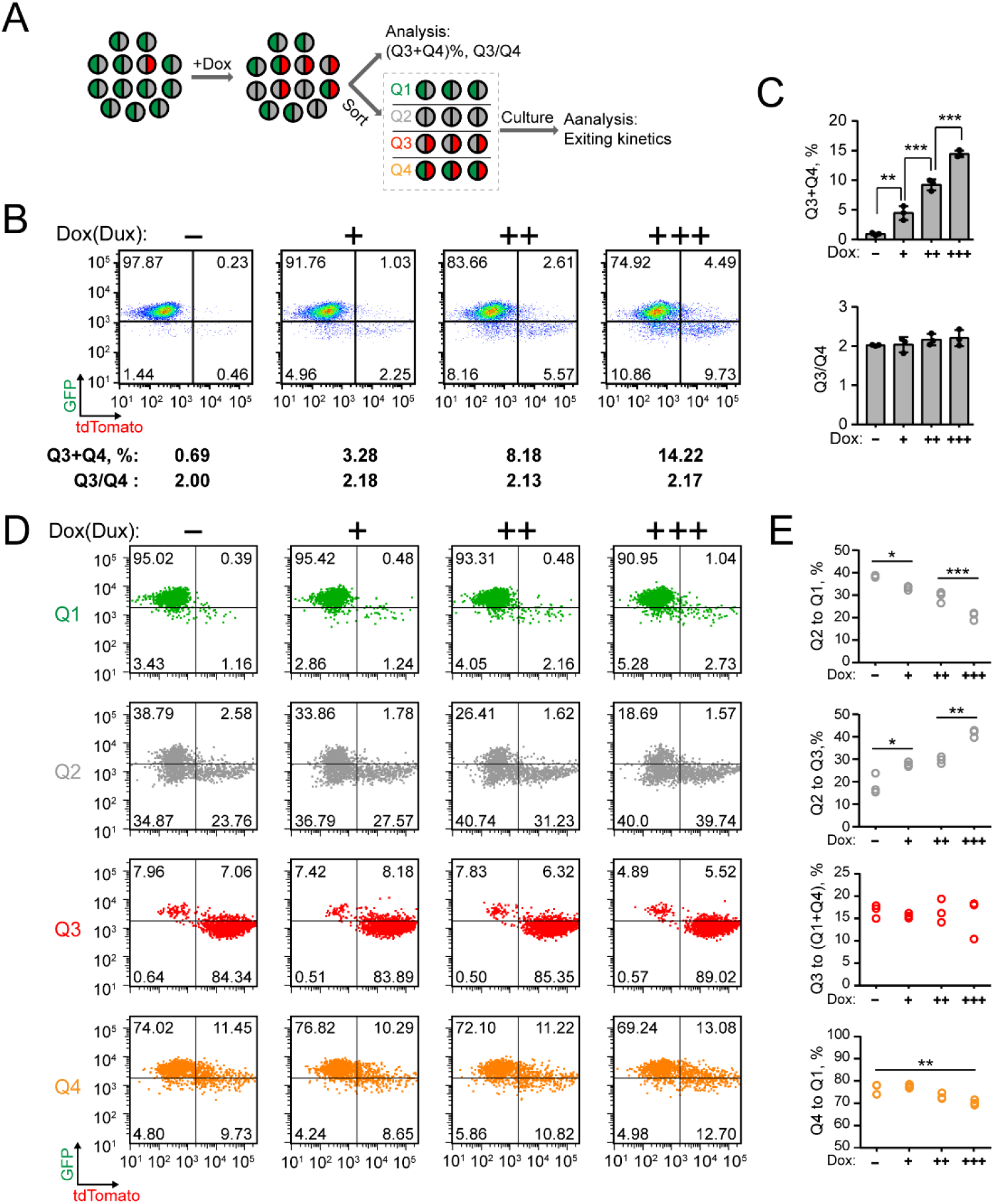
Transient Dux induction promotes the entry of 2C-like state but has little effect on the exit of 2C-like state. Related to Figure 7. (A) Experimental design. (B and C) Flow cytometry analysis of 2C::tdTomato/OCT4-GFP reporter ESCs upon Dux induction by increased dosage of doxycycline; -: 0 nM, +: 10 nM, ++: 20 nM, +++: 40 nM. Representative dot plot with the percentage of Q3+Q4 and Q3/Q4 ratio indicated below (B). Quantification of the percentage of Q3+Q4 and Q3/Q4 ratio (C). Shown are mean ± SD, n = 3 independent experiments. The *p*-value was calculated by one-way ANOVA followed by two-tailed Tukey’s test. (D and E) Flow cytometry analysis of dynamic transition of sorted Q1/2/3/4 with Dux induction by increased dosage of doxycycline; -: 0 nM, +: 10 nM, ++: 20 nM, +++: 40 nM. Representative dot plot (D); Quantification of the percentage of indicated populations (E). Each circle represents one independent experiment, n = 3. The *p*-value was calculated by one-way ANOVA followed by two-tailed Tukey’s test.

**Figure S7:**
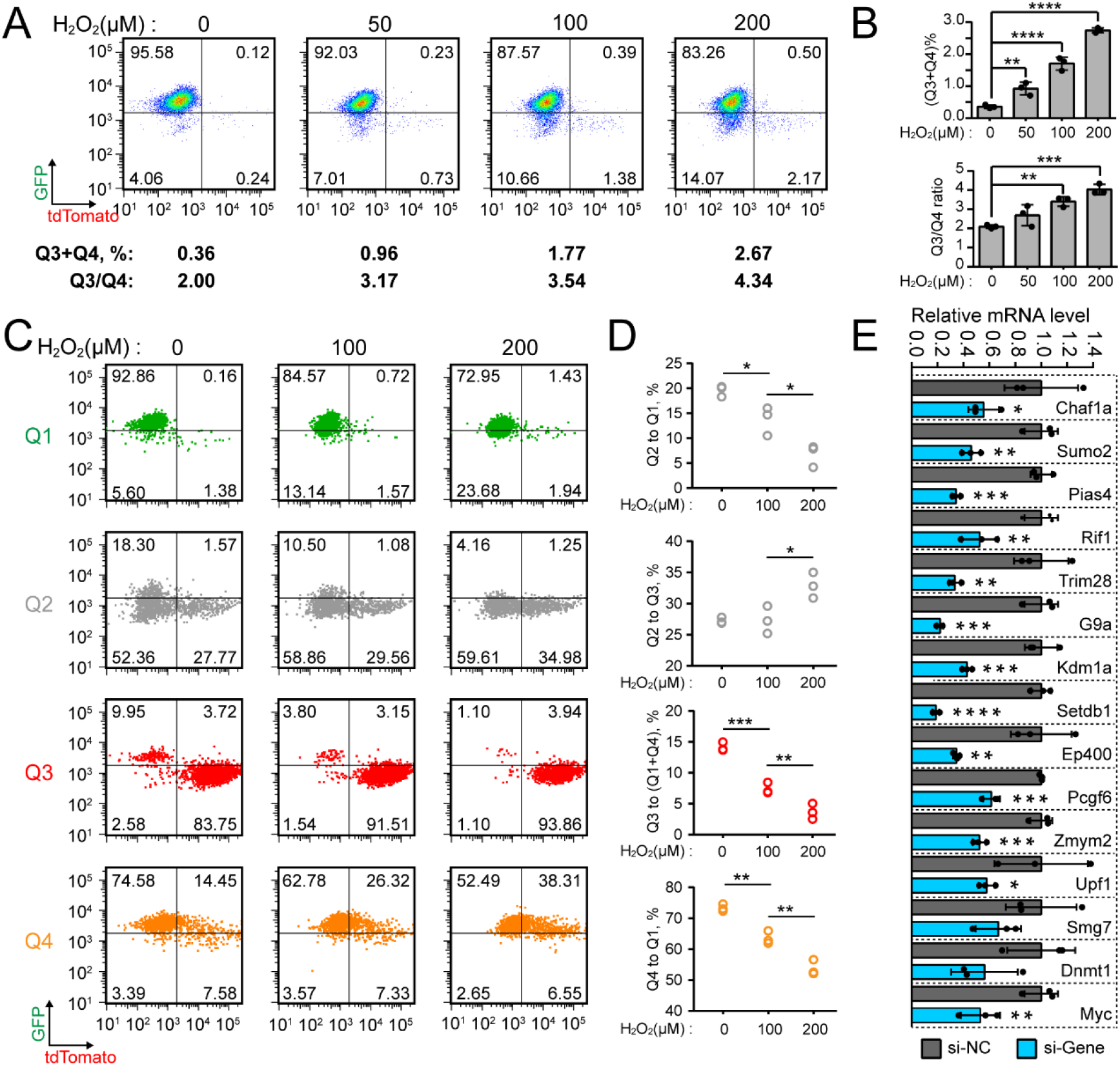
Treatment by H_2_O_2_ promotes the entry of 2C-like state and inhibits the exit of 2C-like state. Related to Figure 7. (A and B) Flow cytometry analysis of 2C::tdTomato/OCT4-GFP reporter ESCs treated with increased dosage of H_2_O_2_. Representative dot plot with the percentage of Q3+Q4 and Q3/Q4 ratio indicated below (A); Quantification of the percentage of Q3+Q4 and Q3/Q4 ratio (B). Shown are mean ± SD, n = 3 independent experiments. The *p*-value was calculated by one-way ANOVA followed by two-tailed Dunnett’s test. (C and D) Flow cytometry analysis of dynamic transition of sorted Q1/2/3/4 treated with increased dosage of H_2_O_2_. Representative dot plot (C); Quantification of the percentage of indicated populations (D). Each circle represents one independent experiment, n = 3. The *p*-value was calculated by one-way ANOVA followed by two-tailed Tukey’ test. (E) RT-qPCR analysis of candidate genes after transfection of corresponding siRNAs against each gene. Shown are mean ± SD, n = 3 independent experiments. The *p*-value was calculated by unpaired one-tailed Student’s *t* test.

**Figure S8:**
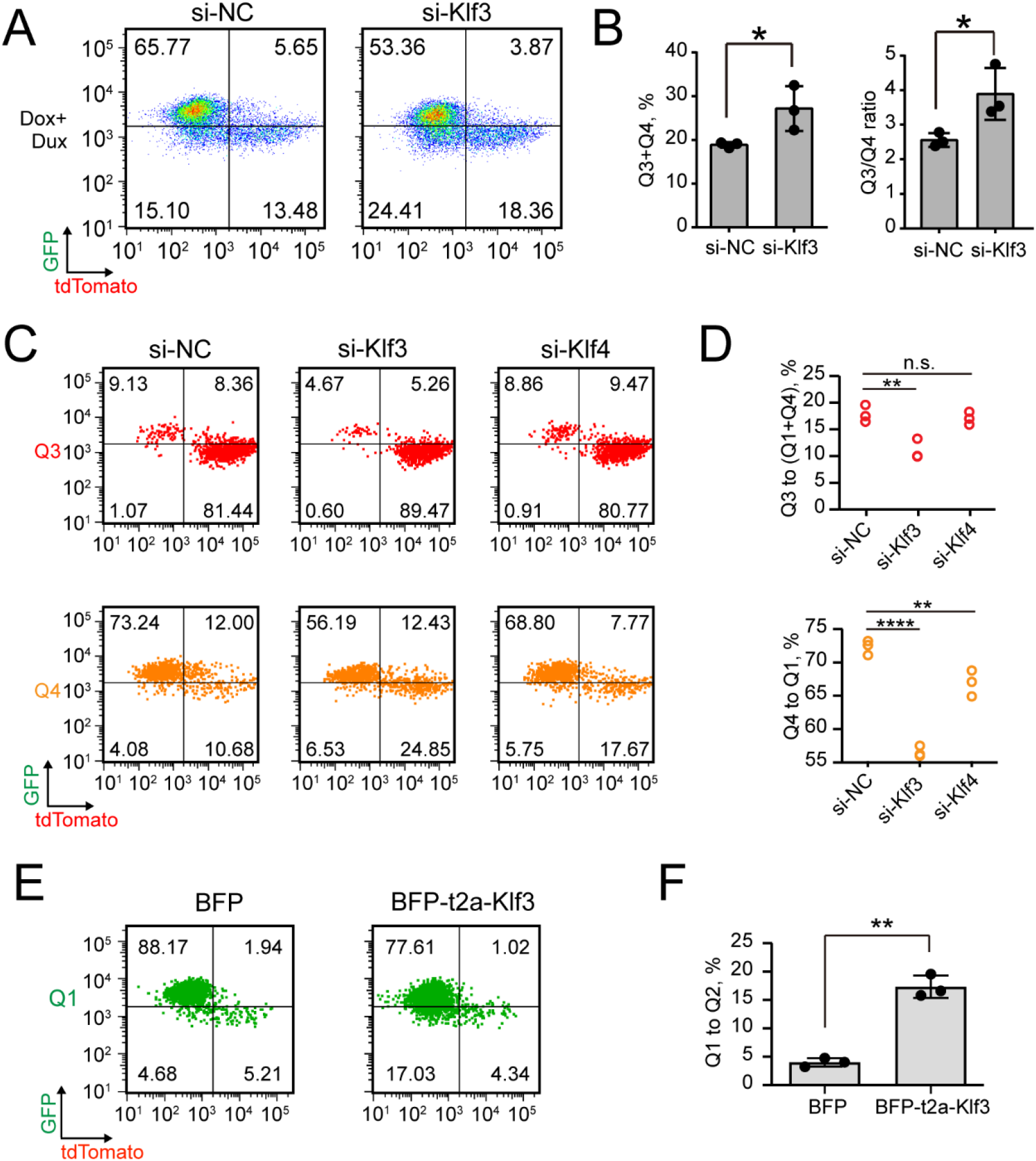
Klf3 is required for efficient 2CLPT. Related to Figure 8. (A and B) Flow cytometry analysis of 2C::tdTomato/OCT4-GFP reporter ESCs treated with si-NC and si-Klf3. 50 nM doxycycline was used to induce Dux expression. Representative dot plots (A); Quantification of the percentage of indicated population (B). Shown are mean ± SD, n = 3 independent experiments. The *p*-value was calculated by unpaired two-tailed Student’s *t* test. (C and D) Flow cytometry analysis of the dynamic transition of sorted Q3 and Q4 pre-treated with si-NC, si-Klf3 and si-Klf4. Representative dot plots (C); Quantification of the percentage of indicated populations (D). Each circle represents one independent experiment, n = 3. The *p*-value was calculated by one-way ANOVA followed by two-tailed Dunnett’s test. (E and F) Flow cytometry analysis of dynamic transition of sorted Q1 transfected with BFP and BFP-t2a-Klf3 mRNAs. Representative dot plots (E); Quantification of the percentage of indicated populations (F). Each circle represents one independent experiment, n = 3. The *p*-value was calculated by paired two-tailed Student’s *t* test.

